# BTK promotes neuroinflammation by interacting with hub genes and modulating microglia following intracerebral hemorrhage

**DOI:** 10.64898/2026.03.04.709243

**Authors:** Siqi Xia, Gao Chen

**Author notes:** **Corresponding authors:** Second Affiliated Hospital, School of Medicine, Zhejiang University 88 Jiefang Road, Hangzhou, 310016, China., E-mail address: SX /, GC.

## Abstract

Bruton’s tyrosine kinase (BTK) has been reported to be important in the inflammatory response in many diseases. However, its role and explicit mechanisms in intracerebral hemorrhage (ICH) remain unclear. Here, we used a mouse ICH model and transcriptomic datasets to explore the effect of BTK on neuroinflammation after ICH. Inhibiting BTK with ibrutinib alleviated ICH-induced neurological deficits and neuroinflammation in mice. After analyzing RNA-sequencing data of ICH and control mice by weighted gene co-expression network analysis (WGCNA) and protein-protein interaction (PPI) analysis, we found that Btk was a hub gene in the green dynamic module. Also, 12 hub genes that closely interacted with BTK were identified in the key gene module, all having a critical role in the inflammatory process. Then, single cell RNA-sequencing data analysis showed that microglia were the immune cells that expressed the most BTK in the mouse brain. After dividing microglia in ICH mice into BTK_high and BTK_low groups, GO/KEGG enrichment analyses of differentially expressed genes (DEGs) between these two microglia groups revealed that most of the top 30 enriched pathways were immune-related. Then, gene set enrichment analysis (GSEA) of the BTK_high and BTK_low microglia showed that the expression levels of four anti-inflammatory and phagocytosis-related pathways were significantly lower in the BTK_high microglia than in the BTK_low microglia. Furthermore, gene set variation analysis (GSVA) demonstrated that multiple immune pathways were expressed differentially between the two microglia groups. Also, six microglia polarization scores were calculated, and the results showed that the BTK_high microglia tend to polarize towards M1 and M2b states, while the BTK_high microglia towards M2 (M2a, M2c) states. Finally, intercellular communication analysis was conducted, and BTK was revealed to promote communication between microglia and other immune cells both at the general level and in specific inflammatory pathways. In conclusion, our study showed that BTK is critical in promoting post-ICH neuroinflammation, at least partly by interacting with Btk-related hub genes and modulating microglia’s immune pathways, polarization, and intercellular communication.

## 1. Introduction

Intracerebral hemorrhage (ICH) is the most common subtype of hemorrhagic stroke. It has extremely high mortality and disability rates, accounting for approximately 15-20% of all stroke cases and 50% of stroke-related deaths ^1^. Based on its pathogenesis, ICH can be divided into primary and secondary ICH. Primary ICH accounts for 78–88% of all ICH cases, mainly caused by chronic hypertension and cerebral amyloid angiopathy, whereas secondary ICH results from various etiologies, such as ruptured aneurysms, vascular malformations, and coagulopathy ^2^. Despite different etiologies and advances in minimally invasive surgical techniques for hematoma evacuation, the prognosis of patients with ICH remains poor, with many patients dying in the early phase of the disease ^3,4^. This is primarily due to the presence of both primary and secondary brain injuries following ICH. Primary brain injury results from mechanical damage caused by the initial bleeding and subsequent hematoma expansion compressing adjacent brain tissue, typically occurring within the first few hours after ICH onset ^5^. Secondary brain injury rapidly ensues, triggered by iron-containing products released from erythrocytes, cytotoxicity, and perihematomal edema, leading to severe neurological deficits and serving as a critical determinant of poor prognosis in patients.

Studies have shown that neuroinflammation is a key mechanism underlying secondary brain injury in ICH. Erythrocytes extravasating from blood vessels undergo lysis and release iron-containing products such as ferrous iron (Fe²⁺), which exerts pronounced neurotoxicity through generating reactive oxygen species ^6^. More importantly, microglia and astrocytes in the perihematomal region become activated, neutrophils infiltrate from the periphery into the central nervous system (CNS), and a series of toxic mediators are released into the surrounding milieu, thereby triggering a robust inflammatory cascade around the hematoma ^7^. This highly damaging neuroinflammatory process has been extensively studied in animal models, where it progresses over hours, days, and even weeks, providing a potential time window for treatment. At the same time, because neuroinflammation exerts a double-edged sword effect on neurological function, redirecting its role toward beneficial outcomes is likely to improve clinical prognosis. In recent years, numerous studies have focused on attenuating the deleterious inflammatory effects of neuroinflammation while promoting its tissue-reparative properties. Several pharmacological agents have been tested in preclinical animal models and even in clinical trials with promising results ^8^. A deeper understanding of the mechanisms of neuroinflammation following ICH will benefit the optimal use of currently available drugs and the development of more effective therapeutic strategies.

Bruton’s tyrosine kinase (BTK) belongs to the cytoplasmic BTK/Tec family of tyrosine kinases. It plays a critical role in the survival, activation, and function of B cells and myeloid cells ^9^. Multiple BTK inhibitors have entered clinical trials for treating leukemias, lymphomas, multiple sclerosis, and autoimmune diseases ^9,10^. Ibrutinib is the first BTK inhibitor approved by the FDA, and can be administered orally for treating certain lymphomas and leukemias. It has strong blood–brain barrier (BBB) permeability; however, it has relatively poor selectivity for BTK and can cause some adverse effects, including thrombocytopenia, minor bleeding, ecchymosis, rash, diarrhea, atrial fibrillation, and opportunistic infections ^11^. Recently, studies have demonstrated that BTK inhibitors exert rapid anti-inflammatory effects in various inflammatory diseases, making them promising therapeutic agents for many disorders ^11,12^. What’s more, inhibiting BTK has been shown to provide anti-inflammatory benefits in several CNS diseases, such as multiple sclerosis, chronic white matter injury, spinal cord injury, and neuromyelitis optica ^13,14^.

However, the role and explicit mechanisms of BTK in neuroinflammation following ICH remain poorly investigated. The current study employs a mouse model of ICH and integrates multiple approaches, including pharmacological intervention, bulk RNA sequencing (RNA-seq), and single-cell RNA sequencing (scRNA-seq), to explore the effect of the BTK inhibitor ibrutinib on ICH-induced neuroinflammation. We found that ibrutinib attenuated ICH-induced neuroinflammation and improved neurological function after ICH. We also identified Btk-related hub genes highly associated with neuroinflammation and ICH using WGCNA. What’s more, transcriptomic analyses revealed that BTK promote neuroinflammation at least partly by modulating microglia functions, including its neuroinflammatory pathways, polarization, and communication with other immune cells. This study is expected to provide new evidence and insights for treating early brain injury following ICH.

## 2. Materials and methods

### 2.1. Animals

Adult male C57BL/6N mice (6–8 weeks, 22–25 g) were purchased from Charles River Laboratories Co., Ltd. (Beijing, China) and housed in individually ventilated cages (IVC) with free access to food and water. The housing environment was maintained at 22±2 °C, with daily temperature fluctuation ≤4 °C, relative humidity of 50%–60%, noise level ≤60 dB, and a 12-hour light/dark cycle to simulate day–night conditions. Before the experiments, all mice were acclimatized to this environment for at least 3 days. All animal experiments followed the National Institutes of Health (NIH) Guide for the Care and Use of Laboratory Animals and were approved by the Institutional Ethics Committee of the Second Affiliated Hospital of Zhejiang University School of Medicine.

### 2.2. ICH model

The mouse ICH model was established by collagenase injection ^15^. Mice were anesthetized by 1% sodium pentobarbital (50 mg/kg, intraperitoneally) and fixed on a stereotactic frame in the prone position, ensuring that the line connecting the bregma and lambda was horizontal. A small burr hole was drilled with an electric cranial drill at 2.5 mm right of bregma. A 10 μL microsyringe (Hamilton, NV, USA) was loaded with 1.2 μL of 0.1 U/μL type VII collagenase (Sigma-Aldrich, MO, USA). And a needle was inserted through the burr hole at a 5° angle toward the left and advanced to a depth of 3.5 mm to reach the right basal ganglia. Subsequently, 0.7 μL of collagenase was injected using a microinjection pump. Body temperature was maintained throughout the surgery. Mice were put on a heating pad to recover postoperatively and were returned to their home cages once fully awake. For the Sham group, all procedures were identical except that no collagenase was injected.

### 2.3. Animal study design

#### Experiment 1

Forty-three mice were randomly divided into six groups to investigate the temporal dynamic changes in BTK expression after ICH (Figure 1): Sham (n=5), ICH 1d (n=9), ICH 3d (n=9), ICH 7d (n=7), ICH 14d (n=8), and ICH 28d (n=5). WB was performed to assess BTK expression in perihematomal or corresponding Sham brain tissue.

**Figure 1.**
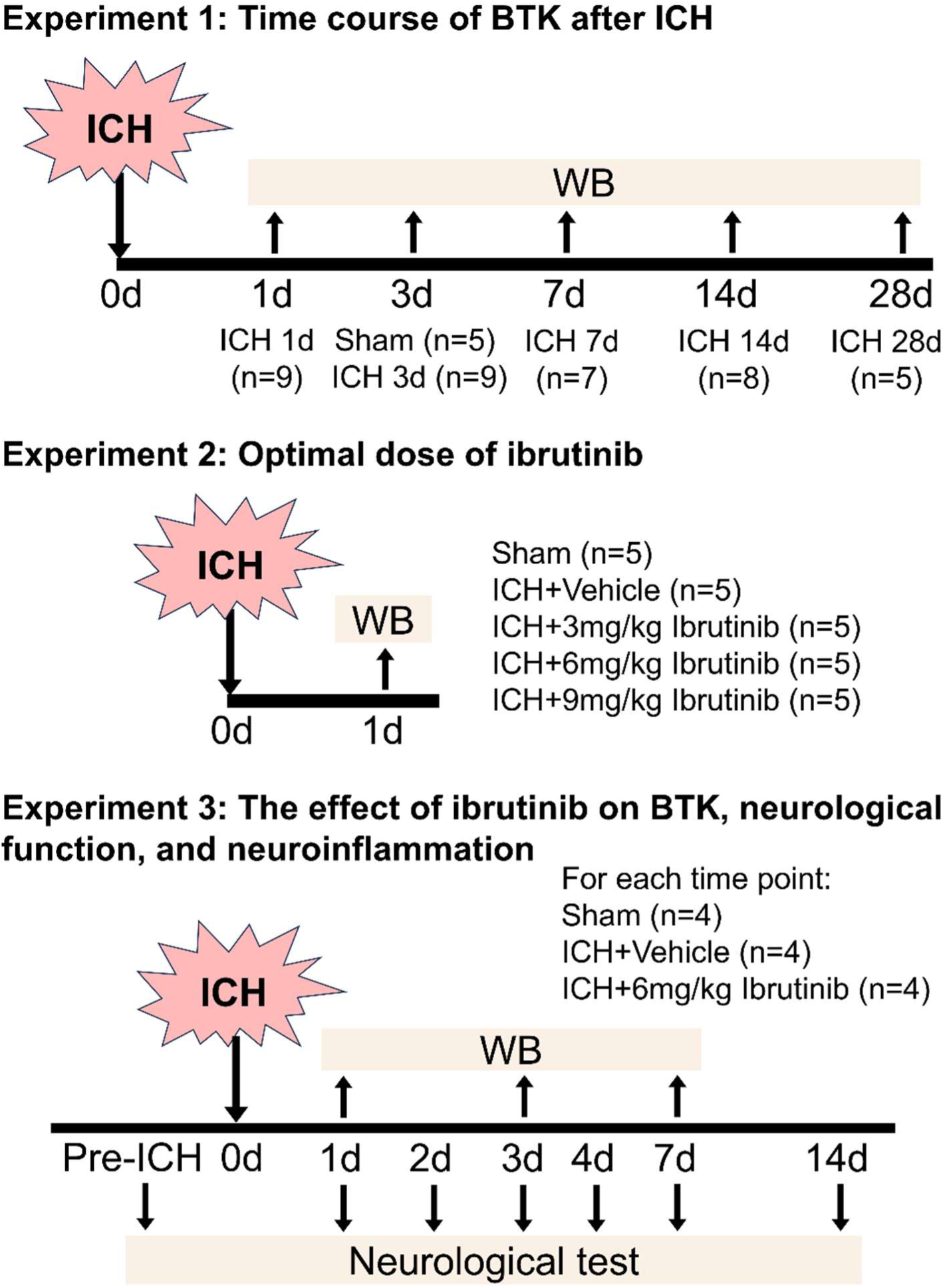
Grouping and experimental design of the animal study.

#### Experiment 2

Twenty-five mice were randomly divided into five groups to determine the optimal dose of ibrutinib for inhibiting BTK in mice (Figure 1): Sham (n=5), ICH+Vehicle (n=5), ICH+3 mg/kg Ibrutinib (n=5), ICH+6 mg/kg Ibrutinib (n=5), and ICH+9 mg/kg Ibrutinib (n=5). At 24 h post-ICH, WB was performed to assess BTK protein expression in perihematomal or corresponding Sham brain tissue.

#### Experiment 3

Forty-eight mice were randomly allocated to three groups to study the role of BTK after ICH (Figure 1): Sham (n=16), ICH+Vehicle (n=16), ICH+6 mg/kg Ibrutinib (n=16). Neurobehavioral tests were performed in each group at pre-ICH and 1, 2, 3, 4, 7, and 14 days after ICH (n=4 per group). To explore the impacts of ibrutinib on BTK expression and neuroinflammation, thirty-six mice were randomly divided into the same three groups (n=12 per group). For each group, WB was used to quantify the protein levels of BTK, inflammatory cytokines, and inflammation-related molecules in perihematomal or corresponding Sham brain tissue at 1, 3, and 7 days post-ICH (n=4 per time point).

### 2.4. Drug administration

The BTK inhibitor ibrutinib (MedChemExpress, NJ, USA) was administered based on the manufacturer’s instructions. A stock solution was prepared by dissolving ibrutinib in DMSO (Sigma-Aldrich, MO, USA) to 55 mg/mL. Before use, the stock solution was diluted with normal saline containing 20% SBE-β-CD (MedChemExpress, NJ, USA) to yield a working solution of 2.75 mg/mL. The vehicle control was prepared by mixing the same proportion of DMSO and 20% SBE-β-CD saline as in the working solution. In the preliminary dose-finding experiment, mice received an intraperitoneal injection of the working solution at graded doses of 3 mg/kg, 6 mg/kg, and 9 mg/kg, or an equivalent volume of vehicle at 30 min after ICH induction to assess lethality and drug effect. Perihematomal or corresponding Sham brain tissue was collected 1 day post-ICH for Western blotting (WB) to measure BTK expression levels. Based on the results (Supplementary Fig. S1), 6 mg/kg was selected as the optimal dose for formal experiments. In the subsequent experiments, mice were administered the first intraperitoneal injection of 6 mg/kg ibrutinib working solution or an equal volume of vehicle at 30 min after ICH, followed by once-daily administration of the same dose per 24 h until the experimental endpoint or postoperative day 7. If a mouse died or the ICH model failed, a backup mouse was used for replacement. The dose gradient used in the preliminary experiment and the post-ICH administration time points were determined based on previous studies ^16,17^.

### 2.5. Neurological function assessment

Four behavioral tests were performed by two researchers who were blinded to the experiment to assess neurological function from 0 to 28 days after ICH, as described in previous literature ^18^. For the corner turn test, each mouse was placed at the entrance of a 30° corner and allowed to walk into the corner before turning either right or left to exit. The direction of each turn was recorded over 10 trials, and the percentage of right turns was calculated. For the forelimb placing test, the mouse was held by the torso with both forelimbs hanging freely. Gently stimulated the whiskers on one side of the mouse with the edge of the cage and observed whether the ipsilateral forelimb grasped the cage rim. Each forelimb was tested 10 times for each mouse, and the percentage of successful placements of the appropriate forelimb onto the countertop edge in response to whisker stimulation was recorded. For the cylinder test, mice were placed individually in a transparent cylinder (8 cm diameter × 25 cm height) and allowed to explore freely until 20 independent rearings were recorded. The initial forelimb (right, left, or both) used to contact the wall during each rear was noted, and a score was calculated as (right-left)/(right + left + both). Higher positive scores indicated greater left-sided hemiparesis. And for the wire hanging test, mice were individually placed on a horizontal stainless-steel bar (50 cm long, 2 mm diameter) placed at 37 cm above a soft-landing surface. Each mouse underwent three 30-second trials. Performance was scored on a 0–5 scale as follows: 0=fell off; 1=clung with both forepaws to the bar; 2=clung with both forepaws and made active attempts to climb onto the bar; 3=clung with both forepaws plus one or both hind paws; 4=gripped the bar with all four paws and wrapped the tail around the bar; 5=escaped to one of the supporting ends.

### 2.6. Western blot

Mice were deeply anesthetized, followed by transcardiac perfusion with ice-cold 0.1 mol/L PBS. The brains were then carefully removed and sectioned into 1-mm-thick coronal slices, which were further dissected to isolate the basal ganglia region containing the hematoma or the corresponding Sham brain tissue. Brain tissue samples were collected and homogenized in cold RIPA lysis buffer (Beyotime, Shanghai, China). Homogenates were centrifuged at 12,000 g for 15 min at 4 °C, and the protein concentration in the supernatant was measured using a BCA Protein Assay Kit (Thermo Fisher Scientific, MA, USA). WB was done as previously described. In brief, 40 μg of total protein from each sample was denatured in loading buffer. Then, the protein was separated by SDS-PAGE, and transferred onto nitrocellulose membranes. After blocking with bovine serum albumin for 1 h, the membranes were incubated overnight at 4 °C with primary antibodies against BTK (1:1000; Cell Signaling Technology, #8547), IL-6 (1:500; Abcam, ab229381), IL-1β (1:500; Abcam, ab234437), TNF-α (1:100; Proteintech, 17590-1-AP), Arg-1 (1:1000; Proteintech, 16001-1-AP), CD68 (1:200; Proteintech, 28058-1-AP), CD11b (1:200; Abcam, ab133357), NLRP3 (1:500; Abcam, ab4207), TLR4 (1:200; Proteintech, 19811-1-AP), and β-actin (1:10000; Proteintech, 66009-1-1g). Membranes were subsequently incubated with horseradish peroxidase-conjugated secondary antibodies (1:10000; ZSGB-Bio, ZB-2301 & ZB-2305) for 1 h at room temperature. Protein bands were visualized using Immobilon ECL Ultra Western HRP Substrate reagent kit (Millipore, MA, USA), and band intensities were quantified by densitometry using ImageJ software (NIH). All experiments had three independent biological replicates per group.

### 2.7. RNA-seq data processing

Original RNA-seq data of three samples in the ICH-3-day group and three samples in the Sham group of the GSE206971 dataset were retrieved from the Gene Expression Omnibus (GEO) database (https://www.ncbi.nlm.nih.gov/geo/) using SRA Toolkit (v.3.2.1) ^19^. The ICH samples were from perihematomal basal ganglia tissues of collagenase injection ICH mouse model and autologous whole blood injection model. The Sham samples were from the same brain region of mice with sham operation. The original data were converted to FASTQ format by FastQC (v.0.12.1) and quality-trimmed by Trimmomatic (v.0.39) to remove low-quality bases ^20^. HISAT2 (v.2.0.4) software was used to map reads to the genome “Mus_musculus.GRCm39”, and the RNA-seq reads of each sample were assembled into final transcripts using StringTie (v.1.3.4-Linux) ^21,22^. Then, the preED.py script in StringTie was used to estimate the expression levels of all transcripts by calculating the raw count of genes. R (v.4.5.1) was used for gene expression data analysis and visualization ^23^. After filtering lowly-expressed genes, differentially expressed genes (DEGs) with |log2(fold change)|˃1 and adjusted p-value˂0.05 were identified using the DESeq2 (v.1.48.2) R package ^24^. The volcano plot was created by the ggplot2 (v.4.0.0) package ^25^, and the heatmap was generated by pheatmap (v.1.0.13) ^26^.

### 2.8. scRNA-seq data processing

CD45^+^ immune cells sorted from an ICH-3-day brain sample and a control brain sample, each containing 15 mice, were extracted from the scRNA-seq dataset GSE230414. R (v.4.5.1) and Seurat (v3.1.5) were used for downstream analysis and visualization ^27^. The ICH sample includes brain tissues of collagenase injection ICH mouse model and autologous whole blood injection model. The Sham sample includes brain tissues of mice with sham operation.

For initial quality control, cells with <500 or >7500 expressed genes in the ICH sample and cells with <500 or >6000 expressed genes in the control sample were removed; also, cells with >15% mitochondrial gene content in the ICH sample and cells with <7% mitochondrial gene content in the control sample were excluded. The distributions of nFeature_RNA, nCount_RNA, and percent.mt of cells in the ICH and Sham groups before and after the quality control are shown in Supplementary Figure S2. Data were then normalized and log-transformed, and the 2000 genes with the highest variance were identified with the FindVariableFeatures function. Dimensionality reduction and cell clustering were subsequently performed. First, principal component analysis (PCA) was applied to the highly variable genes using the RunPCA function. By comprehensively evaluating multiple criteria—including the heatmap of the top 30 principal components, the inflection point in the ElbowPlot, and other diagnostic methods (Supplementary Fig. S3)—the optimal number of principal components was determined to be 29. Next, to address potential technical variability across samples, batch effect correction was performed using the IntegrateLayers function. Then, clustering analysis was conducted on the integrated data based on cell distribution characteristics. Specifically, the first 29 principal components were selected to construct a k-nearest neighbor (kNN) matrix among cells using the FindNeighbors function. Subsequently, cell clustering was performed using the FindClusters function, with the resolution of 0.6. Finally, UMAP was used to visualize the 21 cell clusters on scatter plots (Supplementary Figure S4).

Next, we combined automated and manual approaches for cell type annotation. First, automatic cell annotation was performed using the SingleR package (v.2.10.0) in R, with two separate runs: one based on the immgen reference dataset (2024-02-26) and the other based on the mouse_rnaseq reference dataset (2024-02-26) ^28^. Then, the FindMarkers function was applied to identify marker genes for each cell cluster. Specifically, genes were required to be expressed in at least 25% of the target cell cluster and in other cells, to be highly expressed in the target cluster, and to have a log2foldchange (log2FC)>0.25. Marker genes were then screened using the criteria of adjusted *P*-value (p_val_adj)<0.05 and log2FC>1. A heatmap was generated for the top 10 enriched marker genes of each cell cluster to observe differences in gene expression among the clusters (Supplementary Fig. S5). The identified marker genes for each cell cluster were compared with cell type marker genes collected from the literature and those recorded in the CellMarker database, ultimately yielding the cell type annotations for each cluster. Finally, to validate the accuracy of the cell annotations, UMAP plots were generated based on classic cell type marker genes selected from the literature (Supplementary Table S1, Figure 6-D), and bubble plots were created for the five most specifically expressed genes in each cell type (Supplementary Table S2, Figure 6-E). Inflammatory score and microglial polarization scores were calculated using the AddModuleScore function in Seurat based on a series of marker genes from established literature (Supplementary Table S3).

### 2.9. GO and KEGG pathway analyses

Gene Ontology (GO) is a database that provides a standardized framework for annotating gene functions across three major categories: Biological Process (BP), Molecular Function (MF), and Cellular Component (CC) ^29^. The Kyoto Encyclopedia of Genes and Genomes (KEGG) database contains systematic functional pathways, such as signaling, disease-related, and metabolic pathways ^30^. By mapping the DEGs to GO terms and KEGG pathways, significantly enriched pathways were identified. Both analyses were performed using the clusterProfiler (v4.16.0) package in R.

### 2.10. WGCNA

Weighted Gene Co-expression Network Analysis (WGCNA) was conducted using the WGCNA package (v1.73) in R. Gene co-expression networks were built; genes were clustered into similar expression patterns; and gene modules were identified. Then, correlations among genes within a module, among modules, and between modules and phenotypes were calculated ^31^. In the current study, the soft-thresholding power (β) was selected within the range of 1–40 using the pickSoftThreshold function from the WGCNA package to achieve approximate scale-free topology (R²≥0.9). The lowest power satisfying this criterion was chosen as the optimal soft-thresholding power. Modules were defined using the following parameters: deepSplit=2, minimum module size=30, and mergeCutHeight=0.25. Using module eigengenes (MEs) that represent the first principal component of each gene module, Pearson correlations were computed between modules and the ICH and Sham traits. Hub genes were subsequently screened within key modules based on values of gene significance (GS), module membership (MM), and intramodular connectivity (kWithin).

### 2.11. PPI network

Protein–protein interaction (PPI) networks were built and visualized by Cytoscape (v.3.10.4) software ^32^. Interaction data were obtained from the WGCNA analysis, and only edges with an adjacency value>0.5 were imported into Cytoscape. The PPI network was built and further analyzed using the built-in plugins and algorithms in Cytoscape, with nodes representing proteins and edges denoting connectivity. This enabled the identification of hub genes within the networks.

### 2.12. GSEA

DEGs were ranked by fold-change values and subjected to gene set enrichment analysis (GSEA) using the clusterProfiler (v4.16.0) R package. By comparing the ranked gene list with predefined GO terms, this analysis identified the most enriched biological pathways associated with the observed expression changes and assigned an enrichment score (ES) for each term.

### 2.13. GSVA

Gene Set Variation Analysis (GSVA) was done with the GSVA (v.2.2.0) package to evaluate the expression levels of predefined GO terms across samples ^33^. The “m5.go.v2025.1.Mm.entrez.gmt” gene set from the GO database was used as the reference.

### 2.14. Cell-cell communication

Cell-cell communication analysis was conducted by the CellChat (v.2.2.0) package to infer and visualize signaling networks between immune cells from scRNA-seq data ^34^. Three embedded databases in CellChat—Secreted Signaling, ECM-Receptor, and Cell-Cell Contact were used in this study. This analysis enabled the identification of ligand-receptor interactions and pathway activities, providing insights into how cells influence one another through secreted factors, extracellular matrix mediators, and direct contacts.

### 2.15. Statistics analysis

Continuous data that met normality and homogeneity of variance were presented as mean±SEM (standard error of the mean); otherwise, data were presented as median plus interquartile range. Normally distributed data with equal variances were compared between two groups using the unpaired Student’s t-test, and among three or more groups using one-way analysis of variance (ANOVA) followed by Tukey’s post-hoc test. Data with non-normal distributions or unequal variances were analyzed using the Mann–Whitney U test for two groups or the Kruskal–Wallis test followed by Dunn’s multiple comparison test for three or more groups. A p-value<0.05 indicated statistical significance. All analyses were performed using GraphPad Prism software.

## 3. Results

### 3.1. BTK expression increased in the brain after ICH

RNA-seq data analysis identified 1470 DEGs between the ICH and Sham groups, including 1385 upregulated and 85 downregulated genes (Figure 2-A, B). The GO and KEGG enrichment analyses of these DEGs show that most of the top 30 enriched pathways were related to immune response (Figure 2-C, D), indicating neuroinflammation as a prominent process after ICH, in line with previous studies. Btk was a DEG that significantly increased at 3 days after ICH (log_2_FC=2.35, adjusted p value=7.37*10^-6^) (Figure 2-E). Then, we assessed BTK expression levels in the brain at different time points after ICH. In the ICH mouse model, BTK protein levels significantly increased at 1 day, peaked at 3 days after ICH, then began to decline, and at 28 days after ICH, were not significantly different from normal (Figure 2-F, G, Supplementary Fig. S6).

**Figure 2.**
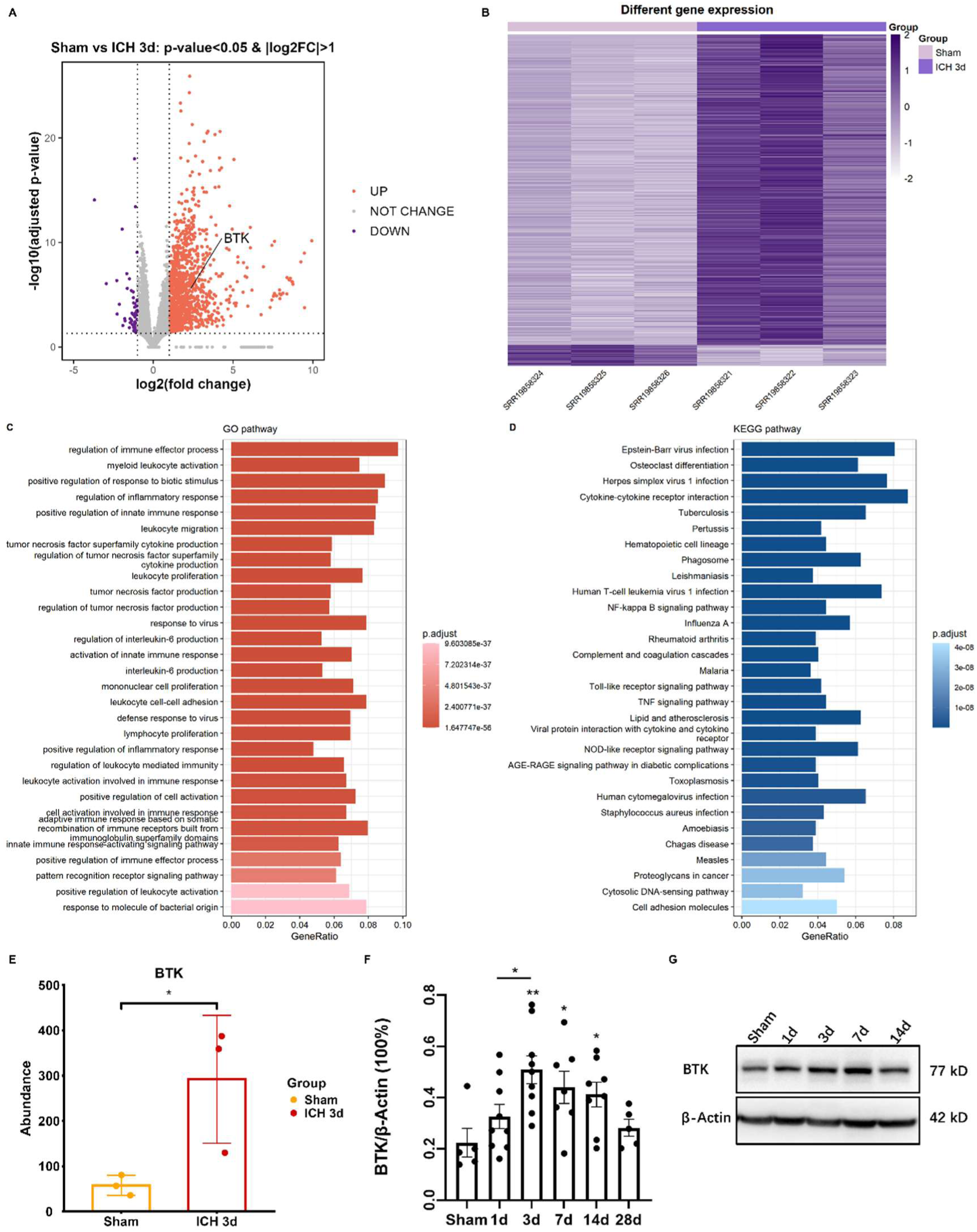
Differentially expressed genes, enrichment analysis, and BTK expression between ICH and Sham. **A-B.** Volcano plot and heatmap of differentially expressed genes (DEGs) between the ICH and Sham groups. **C-D.** Top 30 enriched pathways ordered by adjusted p-value from GO and KEGG enrichment analyses. **E.** BTK transcript levels significantly increased at 3 days after ICH in RNA-seq data. **F-G.** BTK protein levels increased significantly at different time points after ICH in mice. The original blots of Figure G are presented in Supplementary Fig. S6. *P<0.05. **P<0.01 vs. Sham group.

### 3.2. Inhibition of BTK ameliorated ICH-induced neurological deficits

To examine the effect of BTK on ICH-induced neurological deficits, we conducted neurobehavior tests to compare the neurological function among the Sham, ICH+Vehicle, and ICH+Ibrutinib groups. Mice given ibrutinib didn’t present with adverse effects such as thrombocytopenia, minor bleeding, ecchymosis, rash, diarrhea, atrial fibrillation, or opportunistic infections. All groups showed the same baseline of neurological function pre-ICH. In the corner turn test, the ICH+Ibrutinib group showed a significant improvement in neurological function compared with the ICH+Vehicle group on 7 and 14 days after ICH (Figure 3-A). In the forelimb placing test, the ICH+Ibrutinib group had significantly higher functional scores than the ICH+Vehicle group at 1-, 2-, and 14-day post-ICH (Figure 3-B). In the cylinder test, the ICH+Ibrutinib group demonstrated a significant amelioration in neurological deficits compared to the ICH+Vehicle group on days 1, 2, 7, and 14 post-ICH (Figure 3-C). In the wire-hanging test, the ICH+Ibrutinib group exhibited a significant improvement in neurological function compared to the ICH+Vehicle group on 1, 7, and 14 days after ICH (Figure 3-D). Overall, our data indicate that BTK inhibition significantly ameliorated ICH-induced neurological deficits in mice on 1, 2, 7, and 14 days after ICH.

**Figure 3.**
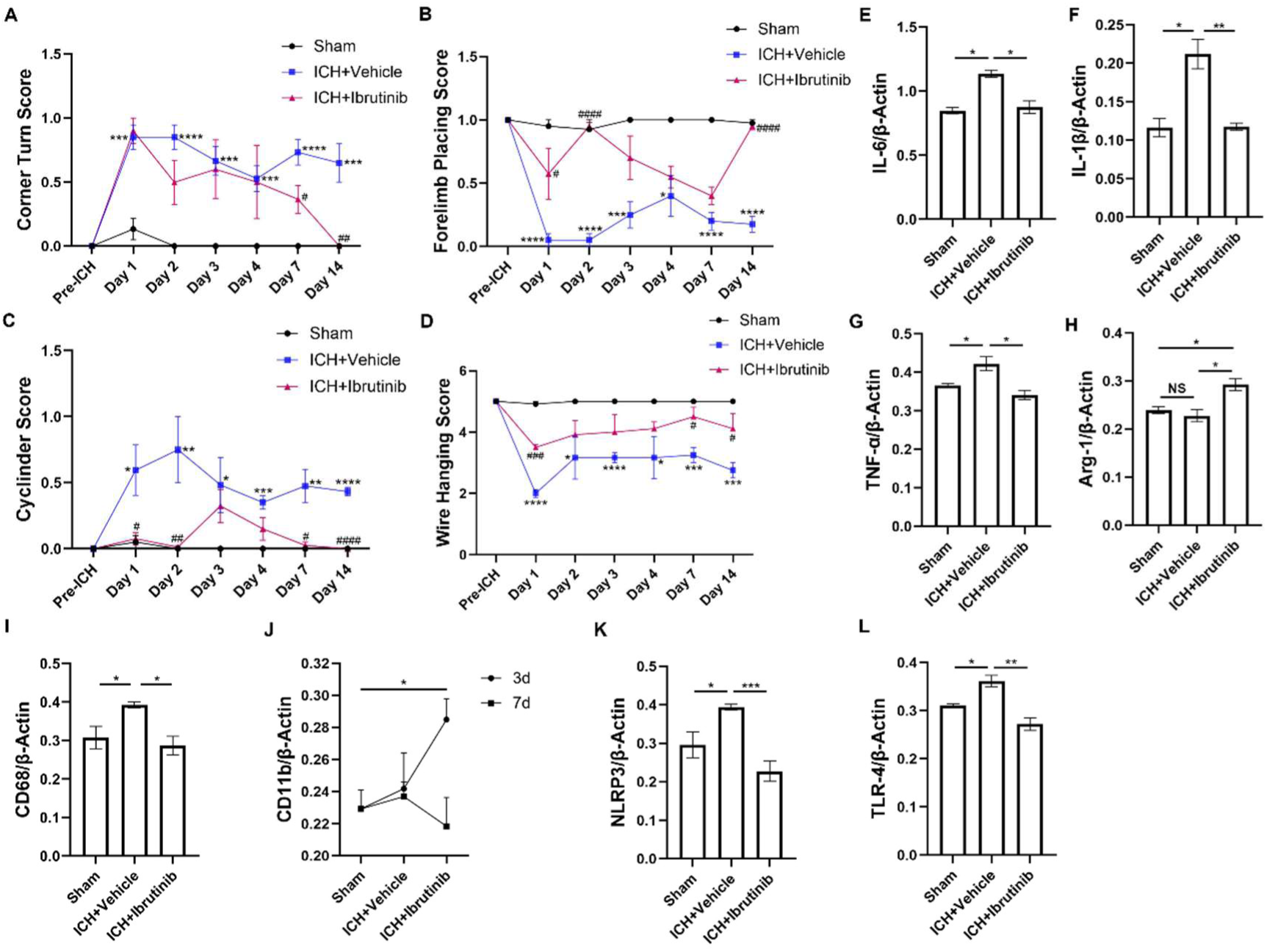
Effects of BTK on neurological function and neuroinflammation after ICH in mice. **A-D.** Corner turn test, forelimb place test, cylinder test, and wire hanging test at 0, 1, 2, 3, 4, 7, and 14 days after ICH (n=4). *P<0.05. **P<0.01. ***P<0.001. ****P<0.0001 vs. Sham group. #P<0.05. ##P<0.01. ###P<0.001. ####P<0.0001 vs. ICH+Vehicle group. **E.** Expression of IL-6 protein on day 1 post-ICH (n=4). **F.** Expression levels of IL-1β protein on day 3 post-ICH (n=4). **G.** Expression levels of TNF-α protein on day 7 post-ICH (n=4). **H.** Expression levels of Arg-1 protein on day 3 post-ICH (n=4). **I.** Expression levels of CD68 protein on day 1 post-ICH (n=4). **J.** Expression levels of CD11b protein on day 3 and day 7 post-ICH (n=4). **K.** Expression levels of NLRP3 protein on day 1 post-ICH (n=4). **L.** Expression levels of TLR-4 protein on day 1 post-ICH (n=4). E-L: NS for P>0.05. *P<0.05. **P<0.01. ***P<0.001.

### 3.3. Inhibition of BTK alleviated ICH-induced neuroinflammation

To investigate the role of BTK inhibition in neuroinflammation after ICH, we examined the protein levels of neuroinflammatory markers in perihematomal brain tissue. The expression levels of proinflammatory cytokine IL-6 on day 1 after ICH (Figure 3-E), IL-1β on day 3 after ICH (Figure 3-F), and TNF-α on day 7 after ICH (Figure 3-G) were significantly higher compared with the Sham group. However, these changes were reversed upon treatment with ibrutinib. Meanwhile, the expression levels of anti-inflammatory cytokine Arg-1 didn’t change markedly on day 3 after ICH compared to Sham, but was substantially higher in the ICH+Ibrutinib group compared to the ICH+Vehicle group (Figure 3-H). What’s more, phagocytic receptor CD68 markedly increased on day 1 post-ICH, and decreased notably when using ibrutinib Figure 3-I); CD11b receptor levels were mildly increased on day 3 and day 7 post-ICH, and with ibrutinib application, markedly increased on 3 days post-ICH while mildly decreased on 7 days post-ICH (Figure 3-J). Existing studies report that ibrutinib can regulate NLRP3 and TLR4 signaling to affect the neuroinflammatory process ^35,36^. Therefore, we also examined the expressions of these two molecules after ICH. The expression levels of both NLRP3 and TLR4 increased significantly on day 1 after ICH and were reversed by ibrutinib treatment (Figure 3 K, L). These results indicate that inhibiting BTK with ibrutinib suppressed the expression of pro-inflammatory molecules while promoting anti-inflammatory cytokines, as well as affected diverse phagocytic receptors and the same receptor at different time points post-ICH differently, suggesting the complex role BTK plays in neuroinflammatory processes.

### 3.4. Btk-related hub genes identified by WGCNA and PPI analysis

To identify the molecular regulatory network of neuroinflammation to which Btk belonged, we selected 5000 high-variance genes with the highest median absolute deviation (MAD) values for WGCNA analysis. No sample was excluded after sample clustering (Supplementary Fig. S7). After building a weighted gene co-expression network with β=30 (Figure 4-A), genes were clustered into 12 static modules and 49 dynamic modules (Figure 4-B). Among all static modules, the turquoise module had the highest positive correlation with the ICH phenotype and thus was regarded as the key module (Figure 4-C, D). Because Btk was located in the turquoise static module and green dynamic module, we chose these two modules for subsequent analyses. Correlation analysis of the turquoise static module revealed that the MM and the GS for the ICH phenotype of all module genes were highly positively correlated (Figure 4-F). What’s more, most genes in the module have |GS|>0.6, further corroborating the high correlation between this module and the ICH phenotype. Since heatmaps of the turquoise static module and green dynamic module display quite similar gene expression patterns (Figure 4-E, Supplementary Fig. S8), and the green dynamic module was contained in the turquoise static module, we chose the green dynamic module for further investigation of Btk-related hub genes.

**Figure 4.**
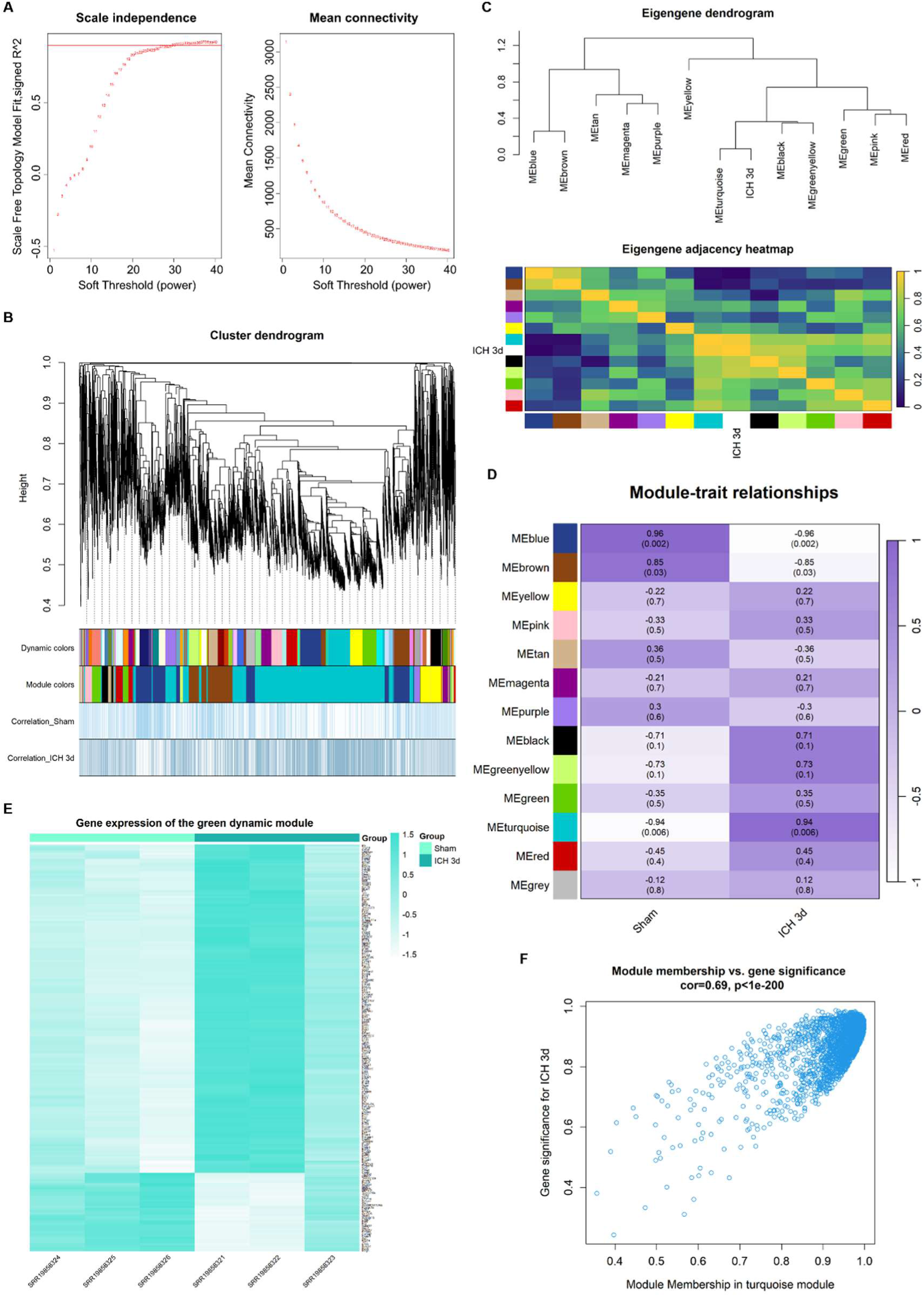
Identification of the key gene module by weighted gene co-expression network analysis. **A.** Soft threshold β=30 was selected based on the combined analysis of scale independence and mean connectivity. The horizontal line of R^2 was set at 0.9. **B.** Co-expression network analysis identified gene modules containing genes with similar expression patterns. Each branch represents a gene, and each color at the bottom represents a gene module. **C.** Cluster dendrogram and adjacency heatmap of all eigengenes and the ICH phenotype. In the heatmap, yellow represents a high adjacency while deep blue represents a low adjacency. **D.** Heatmap of the correlations between modules and the sham and ICH phenotypes, where deep purple indicates a positive correlation, and white indicates a negative correlation. **E.** Heatmap of gene expression in the green dynamic module, where deeper green represents higher expression. **F.** Scatter plot showing correlations between module membership (MM) and gene significance (GS) for the ICH phenotype of all genes in the turquoise static module, where “cor” represents the correlation coefficient between MM and GS.

We found that Btk was the gene with the 10^th^ highest intramodular connectivity (kWithin) in the green dynamic module, indicating it as a critical hub gene of this module. Since all 199 genes in the module had |GS|>0.5, |MM|>0.8, and kWithin>50, we selected 140 genes based on the criteria of p-value(GS)<0.05 and p-value(MM)<0.01. All 140 genes had Btk-related TOM values>0.2, so all were imported into Cytoscape for PPI analysis. First, the MCODE plugin in Cytoscape clustered the genes into two networks—cluster 1 and cluster 2, containing 42 and 27 genes, respectively (Figure 5-A, B). GO analysis of genes in cluster 1, which contained Btk, showed that most of the top 30 enriched pathways were related to cell mitosis (Figure 5-C), suggesting a potential role of BTK in regulating immune cell division and proliferation in neuroinflammation after ICH. Also, the MCC algorithm of the CytoHubba plugin identified 20 hub genes (Figure 5-D). Then, all 140 genes were ranked by Btk-related TOM values, and the top 20 were regarded as highly related to Btk. We found that most of the 20 genes were located in cluster 1 (Figure 5-E). Lastly, convening genes in cluster 1, the 20 hub genes identified by CytoHubba, genes highly related to Btk, and 1470 DEGs between the ICH and Sham group generated 12 Btk-related hub genes, which were Adgre1, S100a10, Emp1, Il10rb, S100a13, Gm13167, Kif22, Sgpl1, Gm7665, Plxnb2, Birc5, and Gas2l3 (Figure 5-F). The detailed information of these hub genes is presented in Table 1. Most of them play important roles in immune processes, such as inflammation, signal transduction, cell migration, and cell division. The PPI network of Btk and these 12 genes is exhibited in Fig. 5G. Therefore, it can be concluded that Btk and the 12 hub genes closely interact to form a regulatory network that modulates neuroinflammation after ICH.

**Figure 5.**
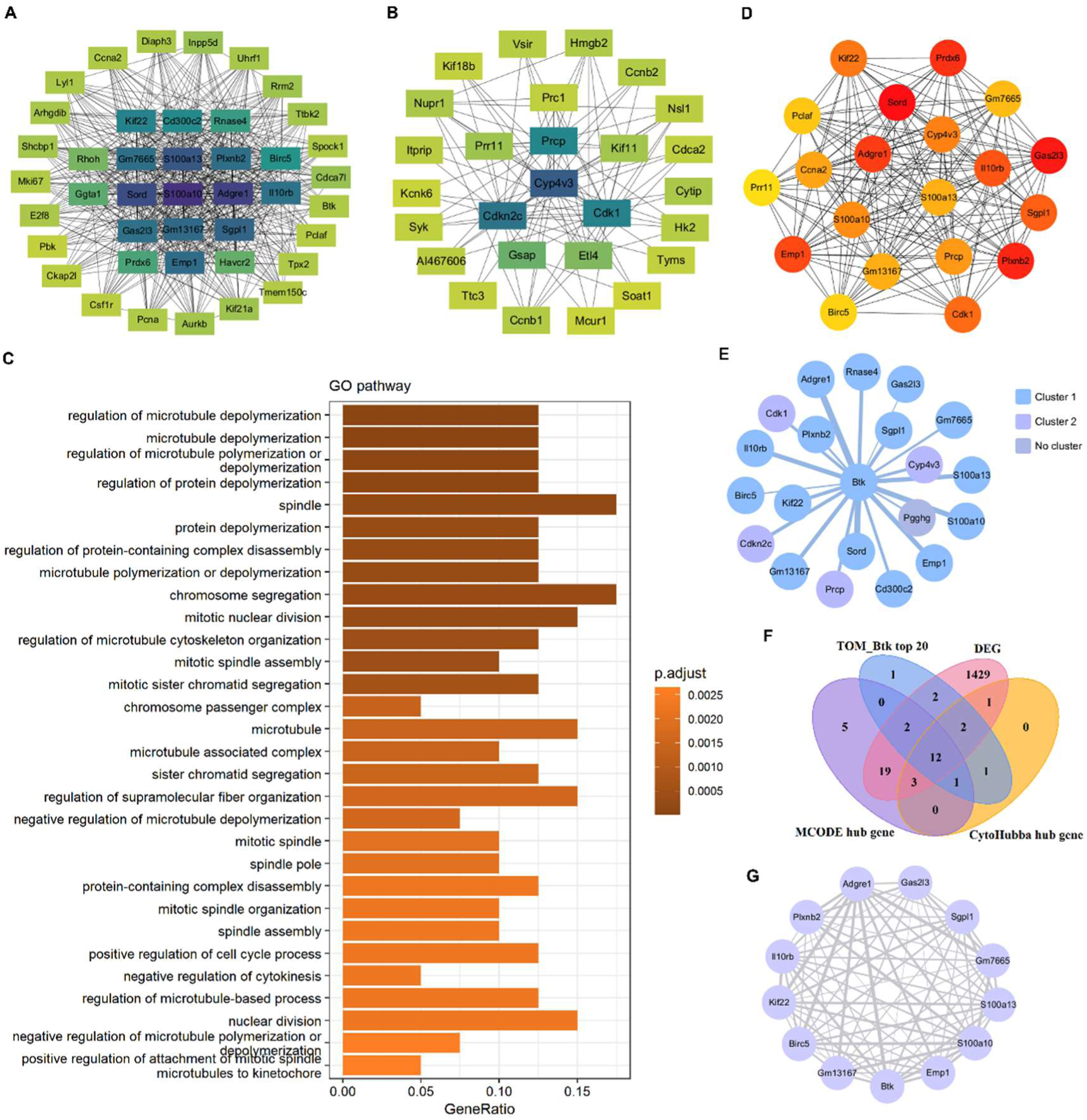
Identification of Btk-related hub genes by protein-protein analysis. **A-B.** PPI networks for cluster 1 and cluster 2 identified by the MCODE plugin in Cytoscape. Darker colors of the squares represent higher connectivity of the gene. **C.** GO enrichment analysis of genes in cluster 1 showed that most of the top 30 enriched pathways were related to cell mitosis. **D.** PPI network containing 20 hub genes identified by the MCC algorithm of the CytoHubba plugin in Cytoscape. Darker colors of the nodes represent higher connectivity of the gene. **E.** Distribution of 20 Btk highly related genes in cluster 1 and cluster 2. Thicker lines represent greater interaction between two genes. **F.** Venn diagram showing the 12 Btk-related hub genes. **G.** PPI network of Btk and Btk-related hub genes. Thicker lines represent greater interaction between two genes.

**Table 1.**
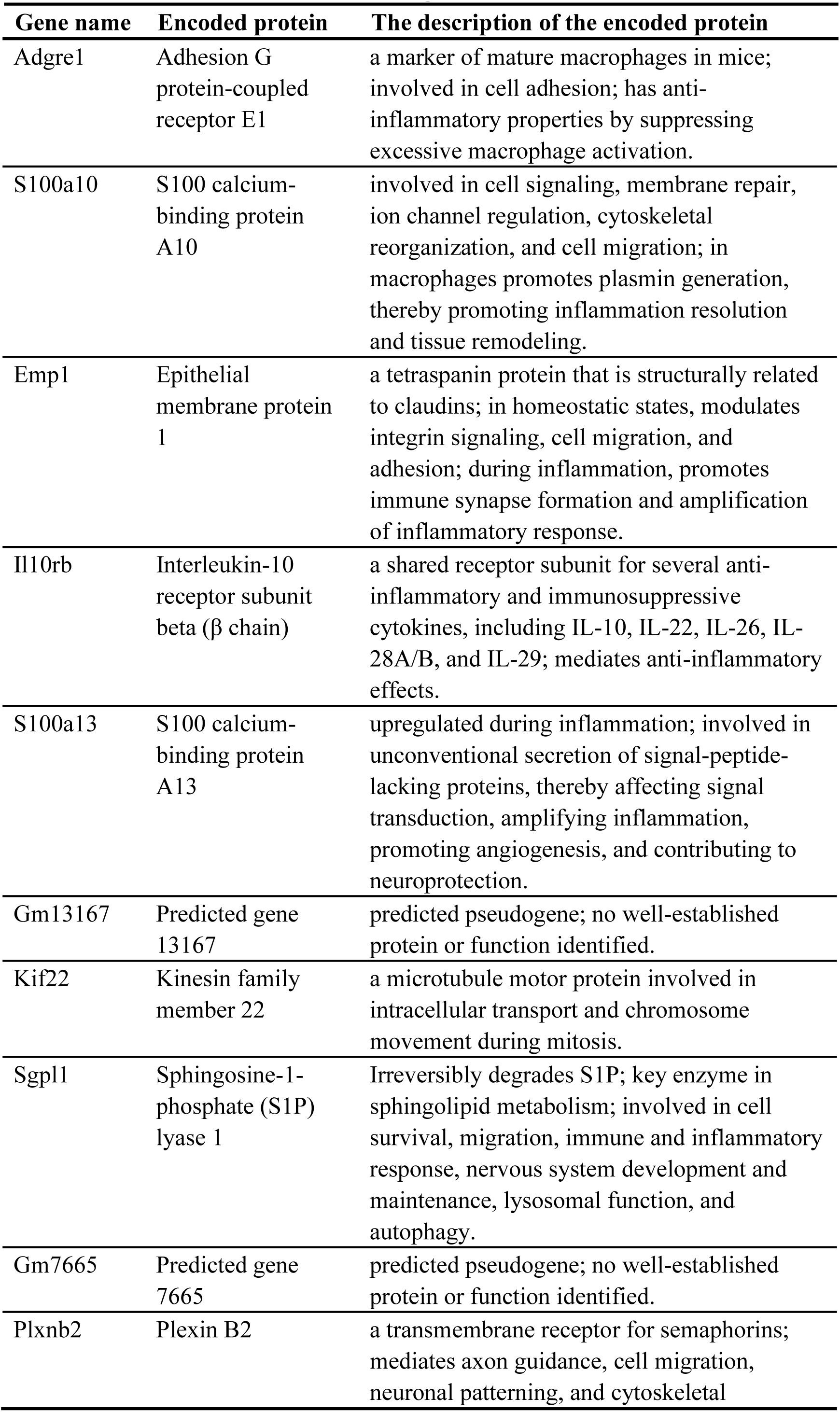
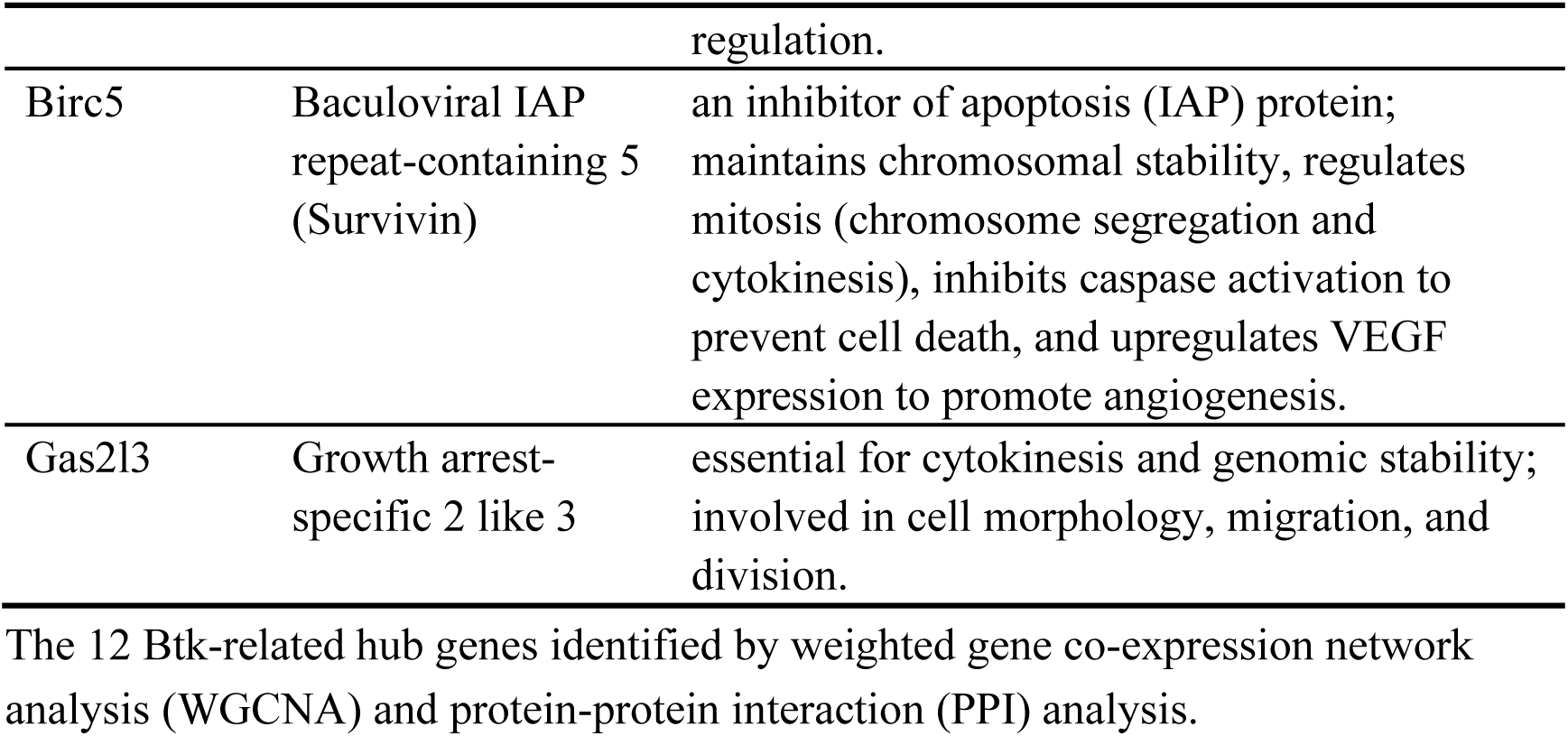
Information of Btk-related hub genes.

### 3.5. BTK modulates neuroinflammatory pathways in microglia

To delineate the cellular landscape of BTK function, scRNA-seq data of mice in the ICH and Sham groups were analyzed. A total of 23312 CD45^+^ immune cells were included, divided into 10 cell types based on their distinct marker genes (Figure 6-A, B, D, E). In line with previous literature, microglia made up the largest proportion of immune cells both before and after ICH (Figure 6-C) and are the immune cells with the third highest inflammatory score (Figure 7-A, B). Further analysis of BTK expression revealed that microglia expressed most (76.4% in the Sham group and 83.7% in the ICH group) of the BTK amount (Figure 7-C, D, F) and had the third highest average BTK expression level (Figure 7-E). These results suggest that microglia are the main immune cells that express BTK after ICH. Therefore, we focused on studying the function of BTK in microglia.

**Figure 6.**
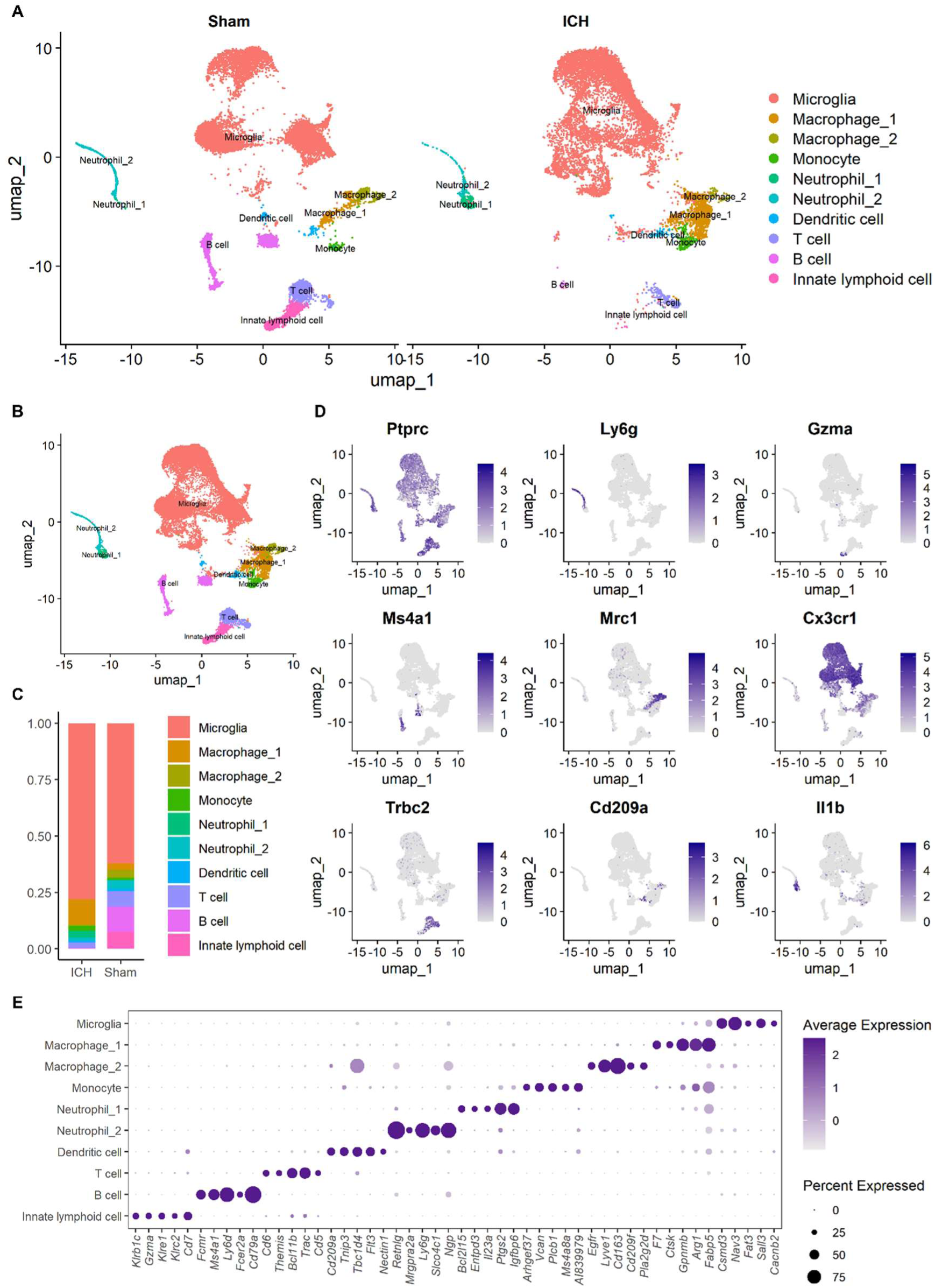
Single-cell analysis of CD45^+^ cells in the brain of ICH mice. **A-B.** UMAP plot of cells from sham (n=10457), ICH (n=12855), and all (n=23312) groups, colored by cell type. **C.** Stacked bar chart showing the proportion of 10 cell types in the sham and ICH groups. **D.** UMAP plots exhibiting selected marker genes from the literature for determining cell identity. **E.** Dot plot showing the average expression (dot color) and percentage of expressing cells (dot size) for the top five marker genes calculated in this study of each cell type.

**Figure 7.**
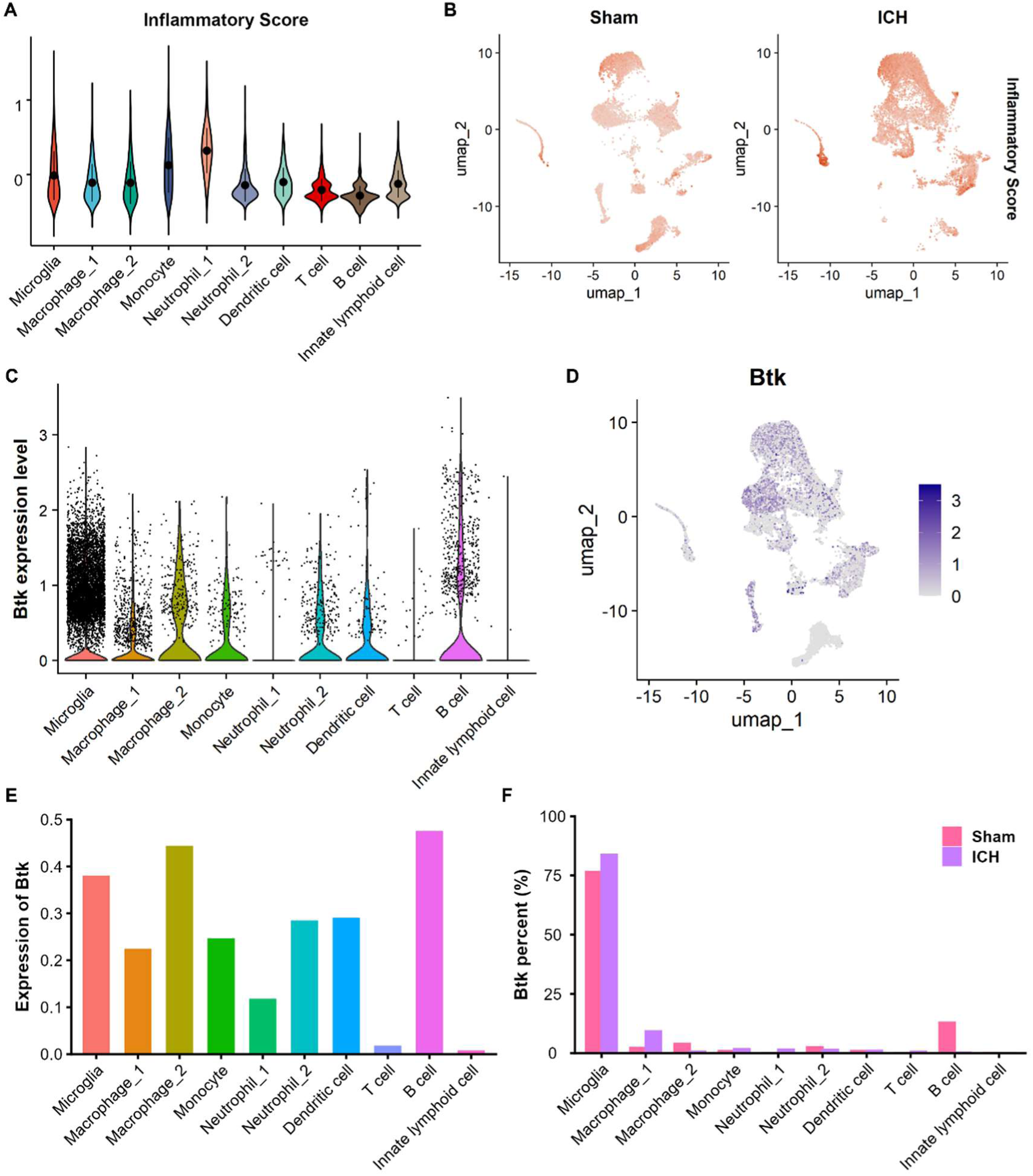
Inflammatory score and BTK expression of different cell types. **A.** Violin plot illustrating inflammatory scores for each cell type in ICH brain, calculated with Seurat’s AddModuleScore function using the inflammatory gene set (Il1a, Il18, Tnf, Tlr2, Tlr4, Nlrp3, Nfkb1, Irf5, Stat1, Socs3, Ccl2, Il6, Ccl3, Ccl4, and Cxcl10). Dots and error bars indicate mean±SD. **B.** UMAP plots showing the distribution of inflammatory scores in sham and ICH groups. Darker yellow represents a higher inflammatory score. **C.** Violin plot illustrating BTK expression levels across cell types, with each dot representing the expression level of one cell. **D.** UMAP plot showing the distribution of BTK expression in all cells. Darker purple represents higher expression. **E.** Bar plot displaying average BTK expression levels in each cell type. **F.** Bar plot showing the percentage of total BTK expression by each cell type in the sham and ICH groups.

Based on the mean BTK expression level (0.2462806) of all microglia in the ICH group, we divided ICH microglia into two subgroups—BTK_high MG and BTK_low MG (Figure 8-A), containing 2332 and 5826 microglia, respectively, and analyzed their differences in the neuroinflammation process. Differential gene expression analysis identified 1998 DEGs between the groups, including 1752 upregulated and 246 downregulated genes (Figure 8-B), as well as 3892 DEGs between microglia of the ICH and the Sham groups, including 1450 upregulated and 2442 downregulated genes (Figure 8-C). Intersecting these two sets of DEGs yielded 602 common DEGs, including 365 upregulated and 237 downregulated genes (Figure 8-D). The enrichment analysis of the 1998 DEGs showed that 12 of the top 30 enriched GO pathways were related to immune response (Figure 8-G), while 10 also existed in the top 30 enriched GO pathways between ICH and Sham microglia and were all immune-related (Figure 8-E); 20 of the top 30 enriched KEGG pathways were related to immune response (Figure 8-G), while 12 also existed in top 30 enriched KEGG pathways between ICH and Sham microglia and were all immune-related Figure 8-F). The high overlaps between the two sets of DEGs and between the two sets of enriched pathways suggest that BTK plays an important role in neuroinflammation after ICH. Emap plot (Figure 9-A) and treeplot Figure 9-B) of the top 30 enriched pathways showed that they could be clustered into several functional modules, such as inflammatory reaction, cell migration, and cell mitosis, indicating the regulating effects of BTK on these modules.

**Figure 8.**
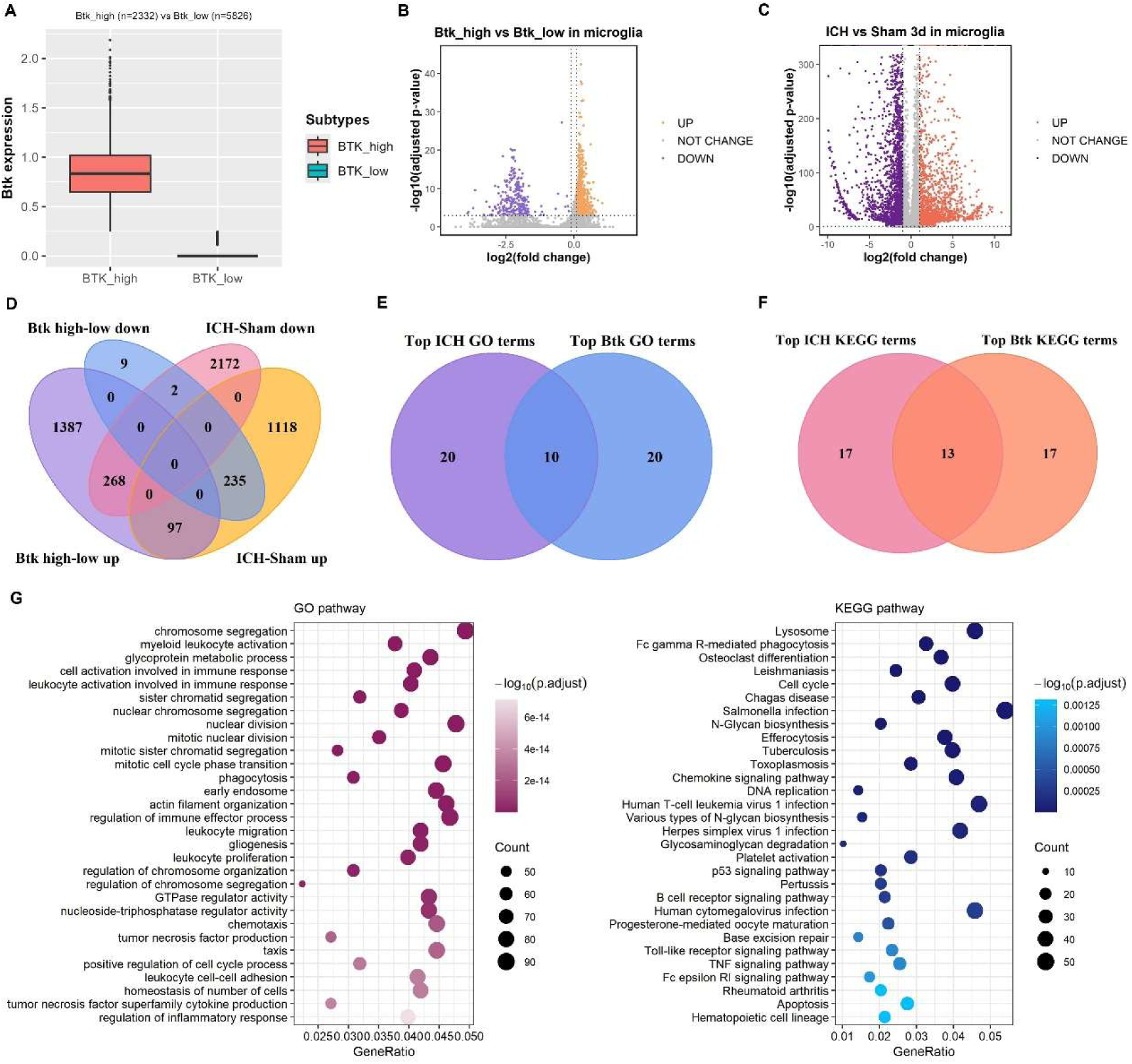
DEGs and enrichment analysis of BTK_high and BTK_low MG in the ICH brain. **A.** Box plot showing BTK expression levels of BTK_high MG and BTK_low MG after ICH. The points above the boxes represent outliers. The upper whisker extends to the maximum value, and the lower whisker extends to the minimum value. The upper edge of the box is the third quartile (Q3), the lower edge is the first quartile (Q1), and the horizontal line inside the box represents the median. The height of the box corresponds to the interquartile range (IQR). **B.** Volcano plot of DEGs between BTK_high MG and BTK_low MG in the ICH brain. **C.** Volcano plot of DEGs between MG in the ICH and in the sham group. **D.** Venn plot exhibiting the intersection of upregulated and downregulated genes in the above two sets of DEGs. **E.** Venn plot showing the intersection of the top 30 GO terms resulting from ICH in microglia and the top 30 GO terms resulting from high BTK expression in microglia. **F.** Venn plot showing the intersection of the top 30 KEGG terms associated with ICH in microglia and the top 30 KEGG terms related to high BTK expression in microglia. G Dot plots exhibiting the top 30 enriched GO terms and KEGG terms resulting from high BTK expression in microglia.

**Figure 9.**
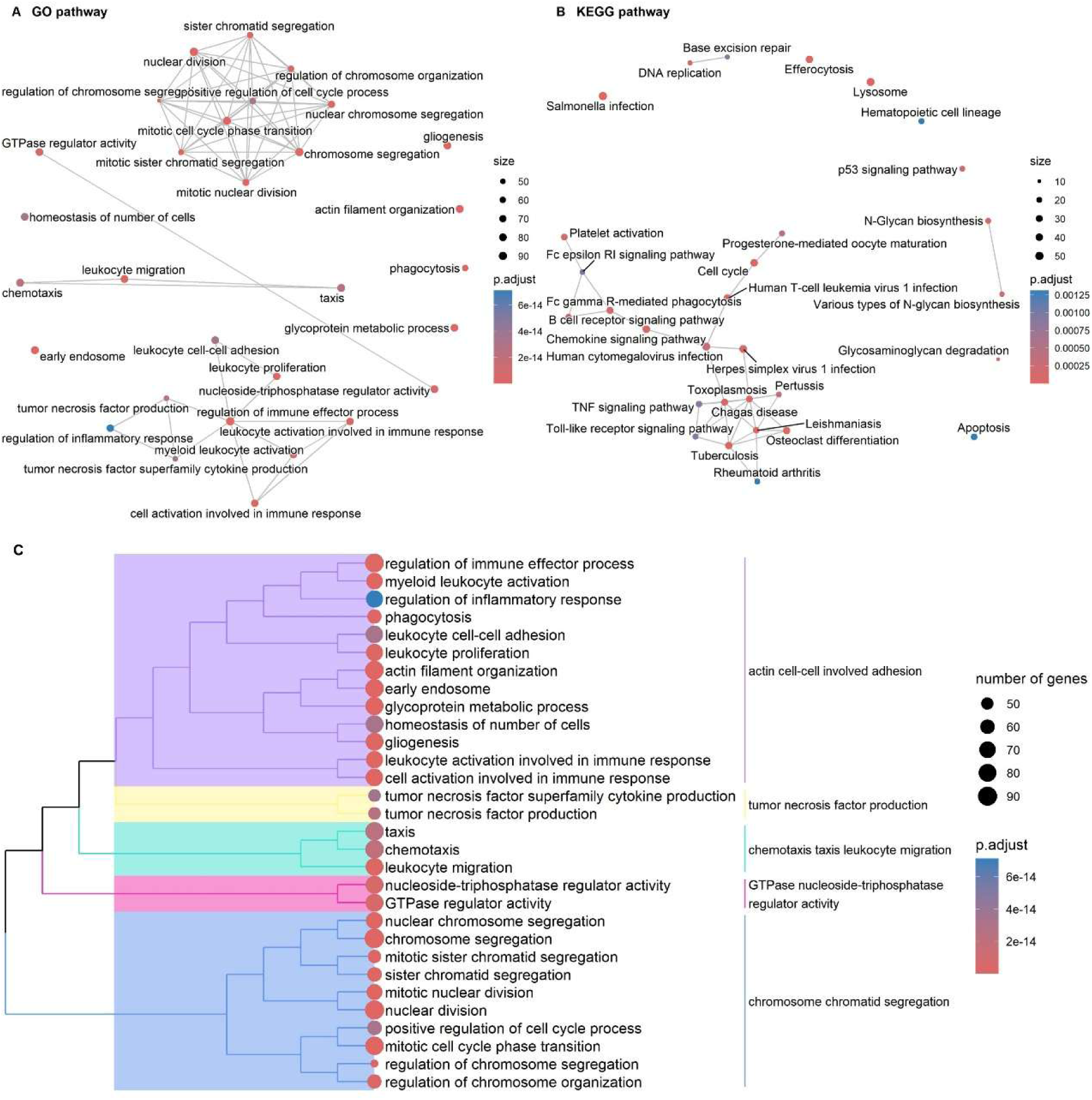
Clustering of the top 30 enriched pathways resulting from high BTK expression in microglia by emap plot and treeplot. **A.** Emap plot showing the top 30 enriched GO pathways resulting from high BTK expression in microglia, clustered into several modules. **B.** Emap plot showing the top 30 enriched KEGG pathways resulting from high BTK expression in microglia, clustered into several modules. **C.** Treeplot exhibiting the clustering of the top 30 enriched GO pathways resulting from high BTK expression in microglia.

GSEA of DEGs between BTK_high MG and BTK_low MG revealed that the TGF-β (Figure 10-A), M-CSF (Figure 10-B), PPAR (Figure 10-C), phagolysosome, and phagosome-lysosome fusion pathways (Figure 10-D) were significantly less enriched in BTK_high MG than in BTK_low MG. GSVA of Sham MG, BTK_high MG, and BTK_low MG showed that BTK_high MG and BTK_low MG had similar levels of glycolytic process through fructose 6-phosphate and of acute phase response, which were higher than Sham MG, suggesting similar cell activities in these two ICH subgroups. Also, compared to both BTK_low MG and Sham MG, BTK_high MG exhibited higher expression of most immune-related pathways such as neuroinflammatory response, innate/adaptive immune response, humoral immune response, microglial cell development/migration/activation/proliferation/differentiation/cytotoxicity, antigen processing and presentation, intracellular receptor signaling, inflammatory apoptosis, pyroptotic inflammatory response, and phagocytosis pathways, while showing lower expression of innate immune response in mucosa and immune memory process pathways (Figure 10-E). Specific to molecular pathways, compared to BTK_low MG and Sham MG, BTK_high MG exhibited higher expression of cell surface PRR, TLR, cGAS-STING, canonical inflammasome assembly, NF-κB, cytokine-mediated signaling, TNF, response to IFN-α/β, GM-CSF, G-CSF, classical complement, and FcR pathways (Figure 10-F); RIG-I, non-canonical inflammasome assembly, IFN-γ, TCR, BCR, and leukotriene pathways were more highly expressed in BTK_high MG than in BTK_low MG, but expression in both groups were lower than in Sham MG (Figure 10-H); TGF-β, M-CSF, PPAR, activin receptor, response to IFN-γ, lectin/alternative complement, netrin, ELRP, ephrin receptor pathways were less expressed in BTK_high MG than in BTK_low MG (Figure 10-G). What’s more, BTK_high MG exhibited lower expression of several growth factor signaling pathways than BTK_low MG (Figure 10-I), including neurotrophin, BDNFR, GDNFR, NGF, FGFR, IGFR, Erbb2-EGFR, VEGF/VEGFR, and PDGFR pathways. These results indicate that BTK upregulates a series of pro-inflammatory pathways while downregulating anti-inflammatory pathways, and suppresses the tissue growth and repair process after ICH.

**Figure 10.**
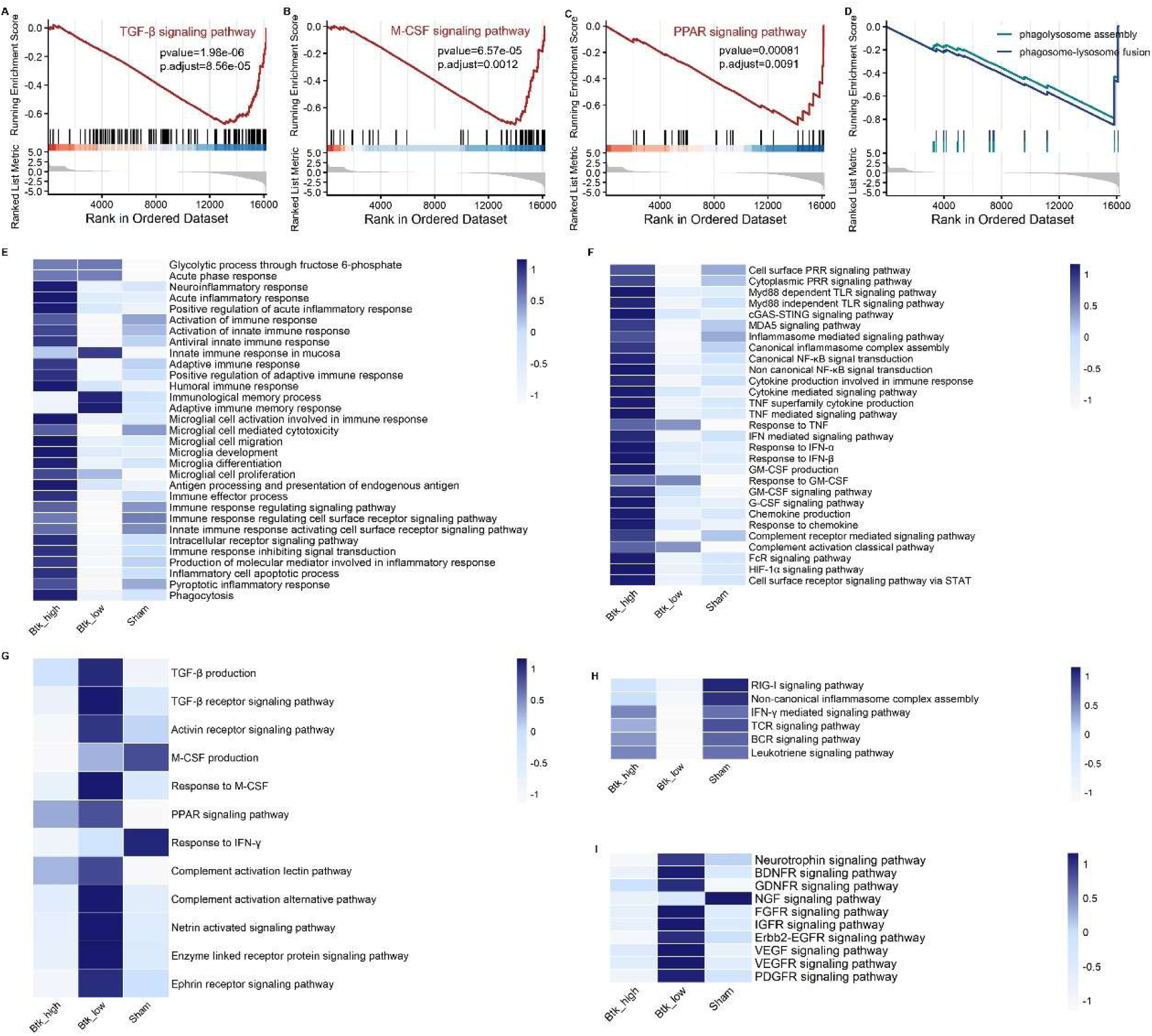
GSEA and GSVA showing differentially expressed inflammatory pathways in BTK_high, BTK_low, and sham MG. **A-D.** GSEA plots showing the TGF-β, M-CSF, PPAR, phagolysosome assembly, and phagosome-lysosome fusion pathways were significantly lower in BTK_high MG compared to BTK_low MG. **E.** Heatmap exhibiting the expression levels of general immune-related gene sets in three groups. **F.** Heatmap exhibiting the levels of inflammatory signaling pathways that are higher in BTK_high MG than in both BTK_low MG and sham MG. **G.** Heatmap displaying the expression of inflammatory signaling pathways that are lower in BTK_high MG than in BTK_low MG. **H.** Heatmap illustrating the levels of inflammatory signaling pathways, which are higher in BTK_high MG than in BTK_low MG but lower in BTK_high MG than in sham MG. **I.** Heatmap exhibiting the expression of growth factor-related signaling pathways in three groups.

### 3.6. BTK affects microglial polarization

Given the vital role of BTK in regulating neuroinflammatory pathways in microglia, we then investigated whether BTK could affect microglial polarization. Scores of six polarization states of microglia were calculated with AddModuleScore and visualized by violin plots. BTK_high MG had significantly higher M1 and M2b scores (Figure 11-A, D) and markedly lower M2 (M2a and M2c) scores (Figure 11-B, C, E) than BTK_low MG. All these scores were higher in the two ICH subgroups than in Sham MG, except for M2c scores, which were similar in BTK_high MG and Sham MG (Figure 11-E). Homeostatic microglia scores were significantly higher in Sham MG than in two ICH subgroups, and were higher in BTK_high MG than in BTK_low MG (Figure 11-F). These results demonstrate that BTK significantly affects microglial polarization, promoting microglia from M2a, M2c, or homeostatic to M1 and M2b phenotypes.

**Figure 11.**
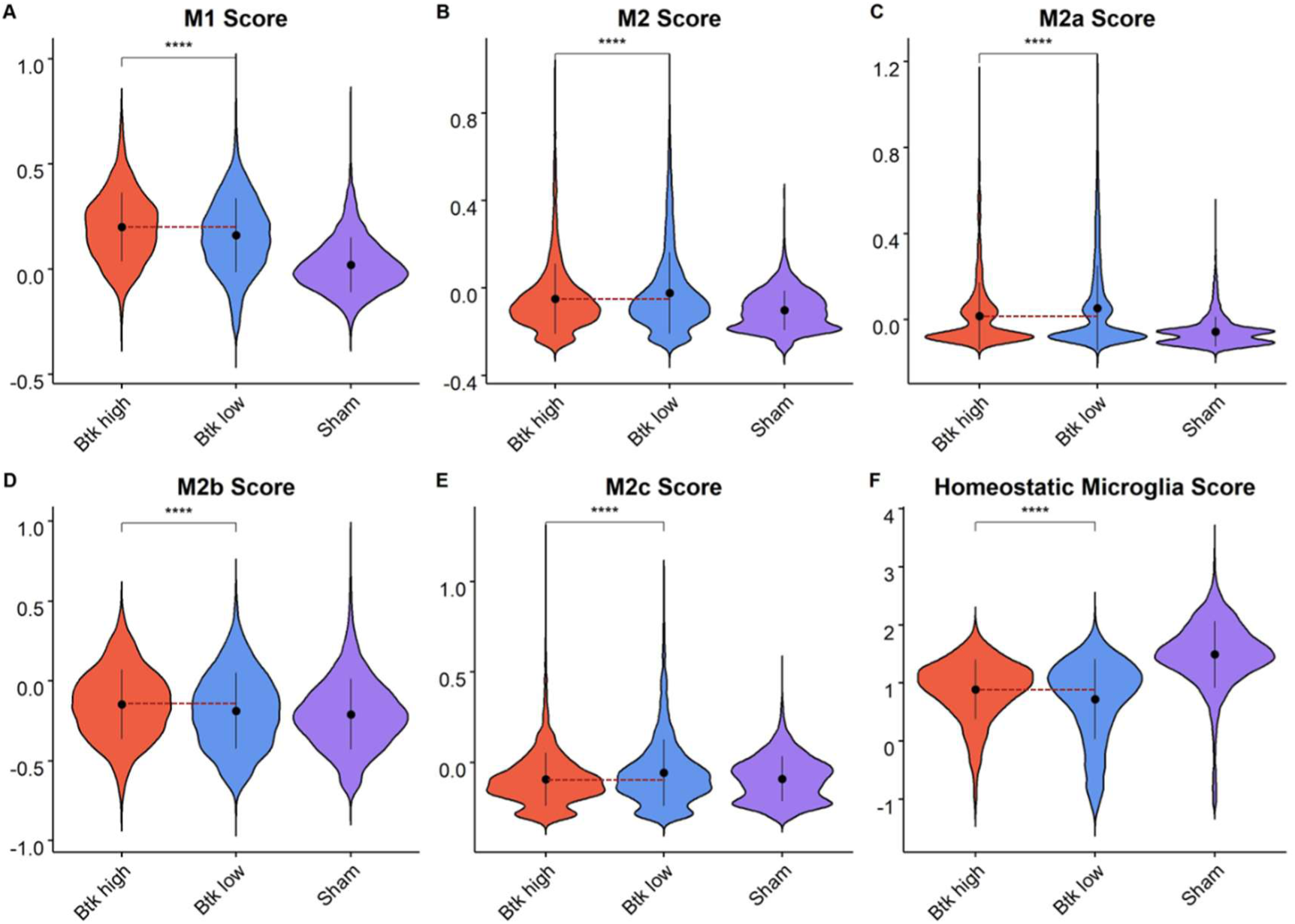
Microglial polarization scores of BTK_high, BTK_low, and sham MG. **A.** Violin plot illustrating M1 scores of microglia calculated by the AddModuleScore function. Dots and error bars indicate mean±SD. **B.** Violin plot showing M2 scores of microglia. **C.** Violin plot exhibiting M2a scores of microglia. **D.** Violin plot displaying M2b scores of microglia. **E.** Violin plot showing M2c scores of microglia. **F.** Violin plot exhibiting homeostatic microglia scores of microglia. ****P<0.0001.

### 3.7. BTK regulates intercellular communication between microglia and other immune cells

Previous studies have reported that crosstalk among immune cells after ICH plays a pivotal role in coordinating neuroinflammation ^8^. Therefore, we used CellChat to analyze the effect of BTK on the intercellular communication between microglia and other immune cells. We found that BTK_high MG had more interactions with all other immune cells than BTK_low MG (Figure 12-A, B, C). Regarding interaction weight, except for a weaker effect on itself and a similar effect on macrophage_1, BTK_high MG exerted stronger effects on any other immune cells than BTK_low MG ((Figure 12-A, B, D). When focusing on specific inflammatory pathways, compared to BTK_low MG, BTK_high MG demonstrated stronger interactions with multiple immune cells in IL-1, IL-2, CXCL, IFN-I, TNF, CCL, CSF, and IL-16 signaling pathways, while having a weaker effect on neutrophil_2 in the CXCL pathway ((Figure 12-E-L). In summary, we conclude that BTK promotes interactions between microglia and most immune cells both at the overall level and in specific inflammatory pathways.

**Figure 12.**
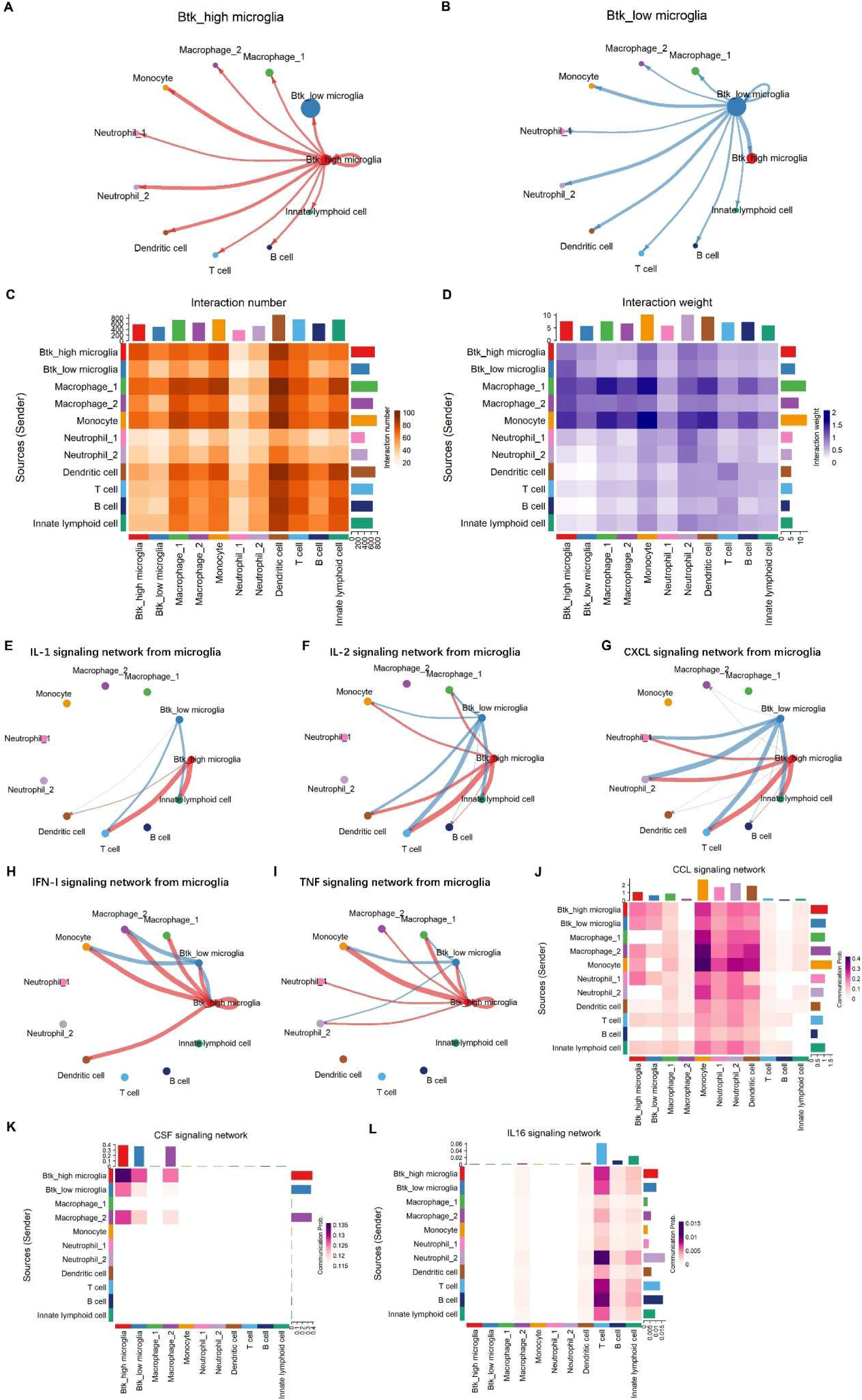
The interactions between immune cells in the mouse brain after ICH. **A-B.** Shell plots exhibiting effects of BTK_high MG and BTK_low MG on immune cells. Arrows indicate the directions of interactions, and thicker lines indicate stronger interactions. **C.** Heatmap showing the interaction numbers between immune cells. Deeper colors indicate more interaction numbers. **D.** Heatmap showing the interaction weights between immune cells. Deeper colors indicate stronger interaction weights. **E-I.** Shell plots illustrating interactions between microglia and immune cells on IL-1, IL-2, CXCL, IFN-I, and TNF signaling pathways. Arrows indicate the directions of interactions, and thicker lines indicate stronger interactions. **J-L.** Heatmaps showing interactions between immune cells in CCL, CSF, and IL-16 signaling pathways. Deeper colors indicate a higher degree of interaction.

## 4. Discussion

In this study, we identified the critical role of BTK in promoting ICH-induced neuroinflammation by interacting with hub genes and modulating microglial functions. First, we found that BTK expression levels were significantly increased after ICH in both RNA-seq data and the mouse ICH model, at the transcriptomic and protein levels, respectively. Second, inhibiting BTK by ibrutinib alleviated neuroinflammation and neurological deficits after ICH. Third, Btk was a hub gene of the green dynamic module, which was highly positively correlated with the ICH phenotype, and interacted closely with hub genes including Adgre1, S100a10, Emp1, Il10rb, S100a13, Gm13167, Kif22, Sgpl1, Gm7665, Plxnb2, Birc5, and Gas2l3. Fourth, among all CD45^+^ immune cells in the ICH brain, microglia expressed the largest proportion of BTK. Fifth, BTK could modulate microglia’s immune response, including upregulating pro-inflammatory pathways, downregulating anti-inflammatory pathways, promoting microglial polarization towards M1 and M2b states, and enhancing intercellular communication between microglia and most other immune cells, both at the overall level and in specific inflammatory pathways. These results demonstrate that BTK is critical in regulating neuroinflammation after ICH, at least in part by interacting with BTK-related hub genes and modulating microglia.

Neuroinflammation is a key mechanism underlying severe secondary brain injury after ICH and is associated with unfavorable clinical outcomes ^4^. Targeting neuroinflammation has been a major research focus for the treatment of ICH ^37^. Numerous preclinical studies and clinical trials have shown that suppressing neuroinflammation after ICH can effectively improve neurological function and clinical outcomes ^8^. BTK belongs to the cytoplasmic BTK/Tec family of tyrosine kinases, which are characterized by the presence of a pleckstrin homology (PH) domain as well as Src homology 3 (SH3) and Src homology 2 (SH2) domains. It plays a crucial role in B-cell receptor (BCR) signaling, as well as in B-cell development, proliferation, and survival. Upon activation of B cells by various ligands, BTK undergoes membrane translocation, a process mediated by its PH domain binding to membrane phosphatidylinositol (3, 4, 5)-trisphosphate [PI(3, 4, 5)P₃]. Once at the plasma membrane, BTK becomes activated through transient phosphorylation of two tyrosine residues, Tyr551 and Tyr223 ^38^. Recent studies have shown that, beyond its roles in B cells, BTK is also involved in multiple immunological signaling pathways in myeloid cells, especially monocytes/macrophages and neutrophils. What’s more, inhibiting BTK exerts anti-inflammatory effects in several CNS diseases ^13,14,38^. For example, a study reported that inhibiting BTK restrained activation of NLRP3 inflammasome, as well as suppressed infarct volume growth and neurological damage in a brain ischemia/reperfusion model in mice, which were likely mediated by inhibiting caspase-1 activation and IL-1β maturation in infiltrating macrophages and neutrophils in the infarcted area of the brain ^36^. Another study showed that ibrutinib reduced LPS-induced neuroinflammatory response in microglia and wild-type mice, probably by inhibiting TLR4 and AKT/STAT3 signaling pathways and prohibiting microglial/astrocytic activation ^16^. In a rat model of SCI, ibrutinib reduced SCI-induced neuroimmune cascade, improved locomotor function, and promoted recovery by inhibiting microglial/astrocytic activation and B cell/antibody response ^39^. In mouse models of stress, BTK inhibition led to anxiolysis and attenuated neuroinflammatory response, as indicated by a significant reduction in NLRP3 inflammasome and IL-1β in the hippocampus and amygdala ^17^. Also, ibrutinib ameliorated neurological impairment, neuroinflammation, BBB disruption, cerebral edema, lipid peroxidation, and neuronal death at 24 h after subarachnoid hemorrhage (SAH) ^40^. However, the specific role and mechanisms of BTK in ICH-induced brain injury remain poorly understood and require further investigation. Therefore, we explored the role of BTK in post-ICH neuroinflammation using a mouse ICH model, together with bulk RNA-seq and scRNA-seq datasets.

First, we found that BTK expression levels were elevated after ICH, peaked approximately 3 days post-ICH, and slowly declined to normal thereafter. Inhibiting BTK markedly alleviated ICH-induced neurological deficits at 1, 2, 7, and 14 days after ICH. What’s more, it decreased the expression of pro-inflammatory cytokines IL-6, IL-1β, and TNF-α in perihematomal brain tissue at 1, 3, and 7 days after ICH, respectively, while increasing the levels of anti-inflammatory cytokine Arg-1 at 3 days post-ICH. This indicates that BTK promotes neuroinflammation after ICH. Meanwhile, ibrutinib significantly downregulated the NLRP3 and TLR4 levels at 1 day post-ICH, corroborating the modulating effects of BTK on these two molecules. Previous studies have reported varying effects of BTK inhibitors on microglial/macrophage phagocytosis. In an in vitro experiment, ibrutinib inhibited antibody-dependent cellular phagocytosis (ADCP) in human monocyte-derived macrophages ^41^. In another study, inhibiting BTK with ibrutinib or siRNA reduced phagocytic function in both mouse microglia and human monocyte-derived macrophages ^42^. Tolebrutinib suppressed microglial phagocytosis in a mouse model of bilateral carotid artery stenosis ^35^. In contrast, evobrutinib enhanced phagocytic capacity in both pro-inflammatory and anti-inflammatory microglia ^43^. Another study showed that fenebrutinib did not affect the ability of human microglia to phagocytose myelin, but ibrutinib at 1 μM significantly reduced this capacity, although the authors attributed this difference to off-target effects of ibrutinib due to its lower selectivity ^44^. In the current study, ibrutinib significantly reduced expression of the phagocytic receptor CD68 at 1 day post-ICH, while markedly increasing expression of CD11b at 3 days post-ICH but slightly decreasing at day 7. Also, GSEA revealed that the expressions of endosome–lysosome assembly and phagosome–lysosome fusion pathways were significantly lower in BTK-high MG than in BTK-low MG; however, GSVA showed that levels of overall phagocytosis-related pathways were significantly higher in BTK-high microglia than in BTK-low MG. Taken together, these findings suggest that BTK and its inhibitors exert complex and context-dependent effects on microglial phagocytosis, likely exerting differential influences on distinct phagocytosis mechanisms, different molecular targets within the same phagocytic pathway, and the same molecule at different time points after ICH.

Next, WGCNA and PPI anlaysis were done to identify BTK-related regulatory network of neuroinflammation. Although WGCNA typically uses a larger sample size (usually n>15) to ensure more reliable gene modules and this study used only six samples, we have taken measures and adjusted parameters carefully to reduce noise and increase statistical power. First, before running WGCNA, we filtered samples based on hierarchical clustering of the sample tree and normalized RNA-seq data using variance-stabilizing transformation (VST) to stabilize variance across low-count genes. Also, we retained only highly variable genes by selecting 5000 genes with top MAD values to focus on those with strong signals. Second, when running WGCNA, we used a high β of 30 with signed R^2>0.9 to reduce spurious connections and set module detection parameters to avoid module oversplitting. We have also run sensitivity analysis by rerunning WGCNA with varied parameters (e.g., different β, deepSplit, and mergeCutHeight) to show that results are not overly sensitive. Therefore, we can ensure a robust correlation network and stable gene modules as manifested in Figure 4-B.

The WGCNA results revealed that Btk was a hub gene of the green dynamic module, which was highly related to the ICH phenotype, and had the 10^th^ highest intramodular connectivity within this module. Most hub genes that interact closely with Btk are highly expressed in immune cells and play critical roles in inflammatory response, signal transduction, cell migration, and cell survival. Adgre1 belongs to the adhesion G-protein-coupled receptors (GPCRs) family. In mice, it is predominantly expressed in mature macrophages and monocytes and is one of the most specific and stable surface markers of macrophages. It participates in cell adhesion and signal transduction, has anti-inflammatory properties by suppressing excessive macrophage activation, and modulates the immune microenvironment in tumor ^45^. S100a1 encodes a member of the S100 family of calcium-binding proteins and serves as a critical bridge between calcium signaling and membrane dynamics. It is highly expressed in immune cells such as monocytes/macrophages, and plays important roles in cell signaling, ion channel regulation, membrane repair, cytoskeletal reorganization, and cell migration. In macrophages, it promotes plasmin generation, thereby contributing to inflammation resolution and tissue remodeling ^46^. Emp1 encodes a tetraspanin protein that is structurally related to claudins. It is mainly involved in signal transduction, epithelial cell differentiation, cell–cell junctions, and tumor progression. In homeostatic conditions, Emp1 is highly expressed in epithelial cells, thereby modulating integrin signaling and regulating cell migration and adhesion; during inflammation, its expression is upregulated in immune cells in response to IFN-γ and TNF-α stimulation, contributing to immune synapse formation and amplification of inflammatory response. Therefore, it is implicated in multiple autoimmune diseases and malignancies ^47^. Il10rb encodes the β chain of the IL-10 receptor and is widely and highly expressed in macrophages, monocytes, lymphocytes, and dendritic cells. It serves as a shared receptor subunit for several anti-inflammatory and immunosuppressive cytokines, including IL-10, IL-26, IL-22, IL-29, and IL-28A/B ^48^. S100a13 encodes an atypical S100 protein that is upregulated during cellular stress, such as inflammation. It facilitates the unconventional secretion of signal-peptide-lacking proteins via a non-ER–Golgi pathway, thereby amplifying inflammation, promoting angiogenesis, and contributing to neuroprotection. It is a key regulator in inflammatory microenvironments and tissue repair processes ^49^. Kif22 encodes a kinesin motor protein primarily involved in intracellular transport and chromosome movement during mitosis. It acts as a crucial motor molecule in cell division ^50^. Sgpl1 encodes sphingosine-1-phosphate (S1P) lyase, which is not only a key rate-limiting enzyme in sphingolipid metabolism that regulates S1P homeostasis, but also an important gene involved in immune and inflammatory response, nervous system development and maintenance, autophagy, and lysosomal function ^51^. Gm13167 and Gm7665 are predicted non-coding RNA genes in the mouse genome with unknown or poorly characterized functions. Plxnb2 encodes a transmembrane receptor highly expressed in the brain. It is a critical component of the semaphorin signaling pathway and plays essential roles in neural development, axon guidance, and cell migration ^52^. Birc5 encodes a core protein inhibitor of the apoptosis process. Its primary functions include maintaining chromosomal stability, regulating mitotic progression, inhibiting apoptosis, and upregulating VEGF expression to promote angiogenesis. It is a well-established cancer biomarker and therapeutic target ^53^. Gas2l3 encodes a protein that modulates cytoskeletal dynamics, cell morphology, migration, and division ^54^. The implications of these hub genes in immune response and the nervous system corroborate the role of BTK in neuroinflammation. Also, GO analysis of genes in cluster 1 showed that the top 30 enriched pathways were mostly related to cell mitosis, suggesting the potential role that BTK may have in immune cell proliferation.

As resident macrophages in the CNS, microglia are the main immune cells that participate in neuroinflammation. They play critical roles in promoting inflammatory response and phagocytic clearance of hematomas, as well as in maintaining homeostasis within the CNS environment ^55^. We found that among all CD45^+^ immune cell types in the brain, microglia had the 3^rd^ highest inflammatory score and the 3^rd^ highest average BTK expression levels, expressing more than 3/4 of the total BTK. Therefore, we focus on studying the function of BTK in microglia. Previous studies have demonstrated that BTK can modulate microglia’s phenotypes and activity and alleviate inflammatory damage. For instance, BTK was reported to participate in the signal transduction pathway that mediates the CCL5-induced elevation in intracellular calcium concentration [Ca(2+)(i)] in human microglia, implicating its potential involvement in inflammatory processes in microglia such as chemotaxis, secretion, and gene expression ^56^. Another study indicated that MOG autoantibodies-induced microglial proliferative response in mice was tightly controlled by Fc receptor and downstream BTK signaling ^57^. In an ischemic demyelination mouse model, a BTK inhibitor suppressed microglial activation and microglia-related inflammation, thus ameliorating white matter injury and cognitive impairments. The authors concluded that the drug probably acted by skewing microglial polarization towards anti-inflammatory and homeostatic phenotypes ^42^. BTK’s role in tissue repair has also been reported. A study indicated that inhibiting BTK promoted remyelination in cerebellar slices and tadpole models, although it could not completely exclude the effect from astrocytes ^58^.

In our study, by analyzing differences between BTK_high MG and BTK_low MG, we found that the DEGs and enriched pathways showed substantial overlap with those between microglia in the Sham and the ICH groups. Also, the top 30 enriched GO and KEGG pathways were highly associated with neuroinflammation. These all suggest the critical role of BTK in modulating neuroinflammation after ICH. Further GSEA analysis revealed that the expressions of three immunomodulatory signaling pathways—TGF-β, M-CSF, and PPAR pathways were significantly lower in BTK_high MG than in BTK_low MG. TGF-β is a classical anti-inflammatory cytokine ^59^. M-CSF is a key cytokine essential for the normal survival, proliferation, and function of microglia. In physiological states, the M-CSF/CSF-1R signaling pathway maintains microglial homeostasis, supports neuronal survival, and promotes the clearance of toxic substances; in disease states, it exerts a dual regulatory effect on neuroinflammation, with the specific outcome depending on the stimulating cytokines, local microenvironment, and stage of the disease. Some studies suggest that it may play an early protective role but become detrimental in later stages ^60–62^. The PPAR pathway exerts potent anti-inflammatory and neuroprotective effects in the CNS by reducing microglial neurotoxicity, inhibiting proinflammatory cytokine production, alleviating oxidative stress, and promoting neuronal differentiation and repair, making it a promising therapeutic target for neuroinflammation-related diseases ^63^. What’s more, GSVA analysis showed that the expressions of multiple proinflammatory pathways were higher while the expressions of inflammation-suppressive and growth factor pathways were lower in BTK_high MG than in BTK_low MG, highlighting the complexity of BTK’s role in neuroinflammation, which includes both pro-inflammatory and anti-reparative effects. Previous studies have divided microglia within the inflammatory microenvironment into different polarization states, including the proinflammatory M1 phenotype, characterized by high expression of cytokines such as TNF-α, IL-1β, and IL-6 and increased neurotoxic potential; and the alternatively activated anti-inflammatory M2 phenotypes (M2a, M2b, M2c), which promote tissue repair, phagocytosis of debris, and secretion of neuroprotective factors such as TGF-β, IL-10, and Arg-1; alongside the quiescent homeostatic state responsible for immune surveillance and synaptic pruning under physiological conditions ^8,64^. In the present study, polarization analysis revealed a significant shift of BTK_high MG towards M1 and M2b phenotypes and a significant shift of BTK_low MG towards M2 (M2a and M2c) phenotypes, while microglia in the Sham group tend to remain homeostatic. These results indicate that BTK critically regulates microglial polarization, favoring M1 and M2b phenotypes after ICH, and suggest that targeted inhibition of BTK may promote transition towards an anti-inflammatory neuroprotective M2 state, thereby alleviating neuroinflammation after ICH. Finally, CellChat analysis of intercellular communication showed that BTK can strengthen communication between microglia and other immune cells both at the overall level and in multiple inflammatory signaling pathways, including IL-1, IL-2, IL-16, IFN-I, TNF, CXCL, CCL, and CSF pathways. All these evidence converge to prove the importance of BTK in microglial activity. Thus, targeting BTK could be an effective way to modulate microglia’s phenotypes and functions and to reduce neuroinflammation after ICH.

While our study demonstrates that inhibition of BTK alleviates neuroinflammation and improves neurological function in a mouse model of ICH, and employs RNA-seq and scRNA-seq data to investigate the underlying mechanisms comprehensively, it also has certain limitations. First, previous studies have shown that, apart from CD45^+^ immune cells, other cell types such as astrocytes and dendritic cells also participate in neuroinflammation and tissue repair following ICH ^59,65,66^. However, the scRNA-seq dataset used in this study included only CD45^+^ immune cells; therefore, the expression of BTK in other cell types and its mechanisms in mediating neuroinflammation require further exploration. Moreover, the animal models, experimental methods, and samples in transcriptomic datasets used in this study are limited. Integration of large-sample, multi-method, and multi-omics datasets in future studies would promote a deeper exploration of BTK’s mechanisms of action and provide a more comprehensive view of its effects. What’s more, the effects of BTK on various phagocytic processes during the inflammatory response, on different molecules within the same phagocytic pathway, the time course of these effects following ICH, and their impact on overall prognosis require further investigation. Lastly, the 12 Btk-related hub genes and multiple immune pathways identified in this study as regulated by BTK need experimental validation in future research.

## 5. Conclusion

Our study shows that inhibiting BTK attenuates neuroinflammation and improves neurological outcomes after ICH, at least partly by interacting with Btk-related hub genes and by modulating microglia’s activity, polarization, and communication with other immune cells. We identify 12 hub genes and multiple signaling pathways associated with BTK, thus providing valuable candidates for future mechanistic and functional investigations. Altogether, the current study supports the notion that targeting BTK might be a promising therapeutic strategy for treating neuroinflammation after ICH.

## Authors contributions

**S. X.** Conceptualization, Methodology, Data curation, Formal analysis, Visualization, Validation, Writing – original draft, Writing – review and editing. **G. C.** Conceptualization, Project administration, Funding acquisition, Resources, Supervision, Writing – review and editing.

## Data availability

The RNA-sequencing data and single-cell RNA-sequencing data are available through Gene Expression Omnibus with accession number GSE206971 and GSE230414, respectively. Other data presented in this study are available from the corresponding author upon request to the corresponding author.

