## Supplementary for "BTK promotes neuroinflammation by interacting with hub genes and modulating microglia following intracerebral hemorrhage"

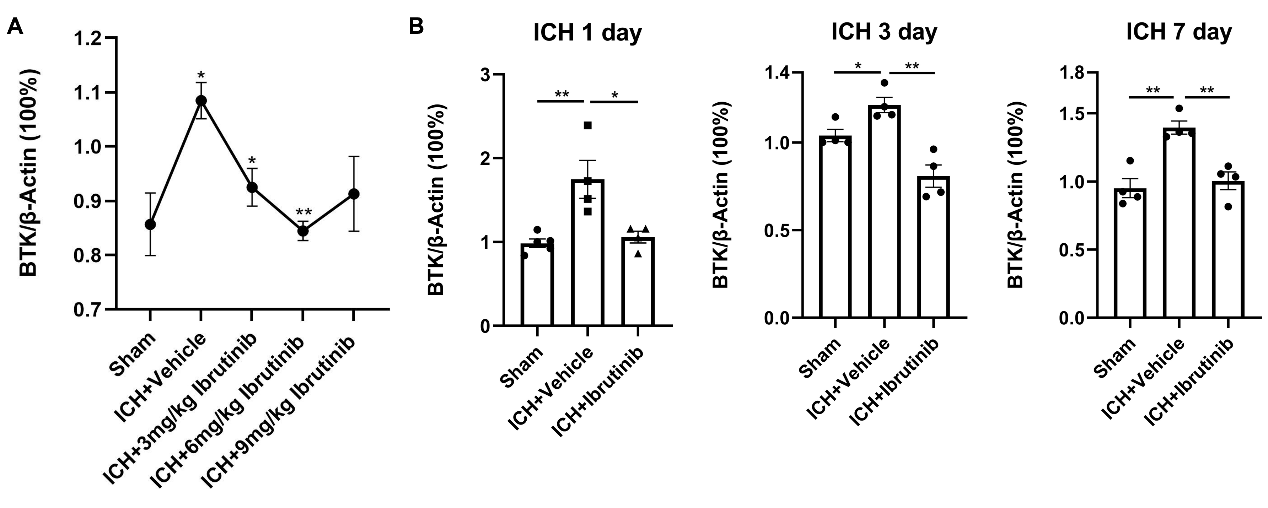
**Supplementary Figure S1.** The inhibition of BTK protein expression by ibrutinib after ICH. **A** The 6 mg/kg dose of ibrutinib exhibited the strongest inhibitory effect on BTK protein expression at 1 day post-ICH, and mice receiving this dose of the drug showed no mortality or adverse effects (n=5). **B** Ibrutinib inhibited BTK protein expression at different time points after ICH (n=4). **P*<0.05. ***P*<0.01 vs. Sham group.

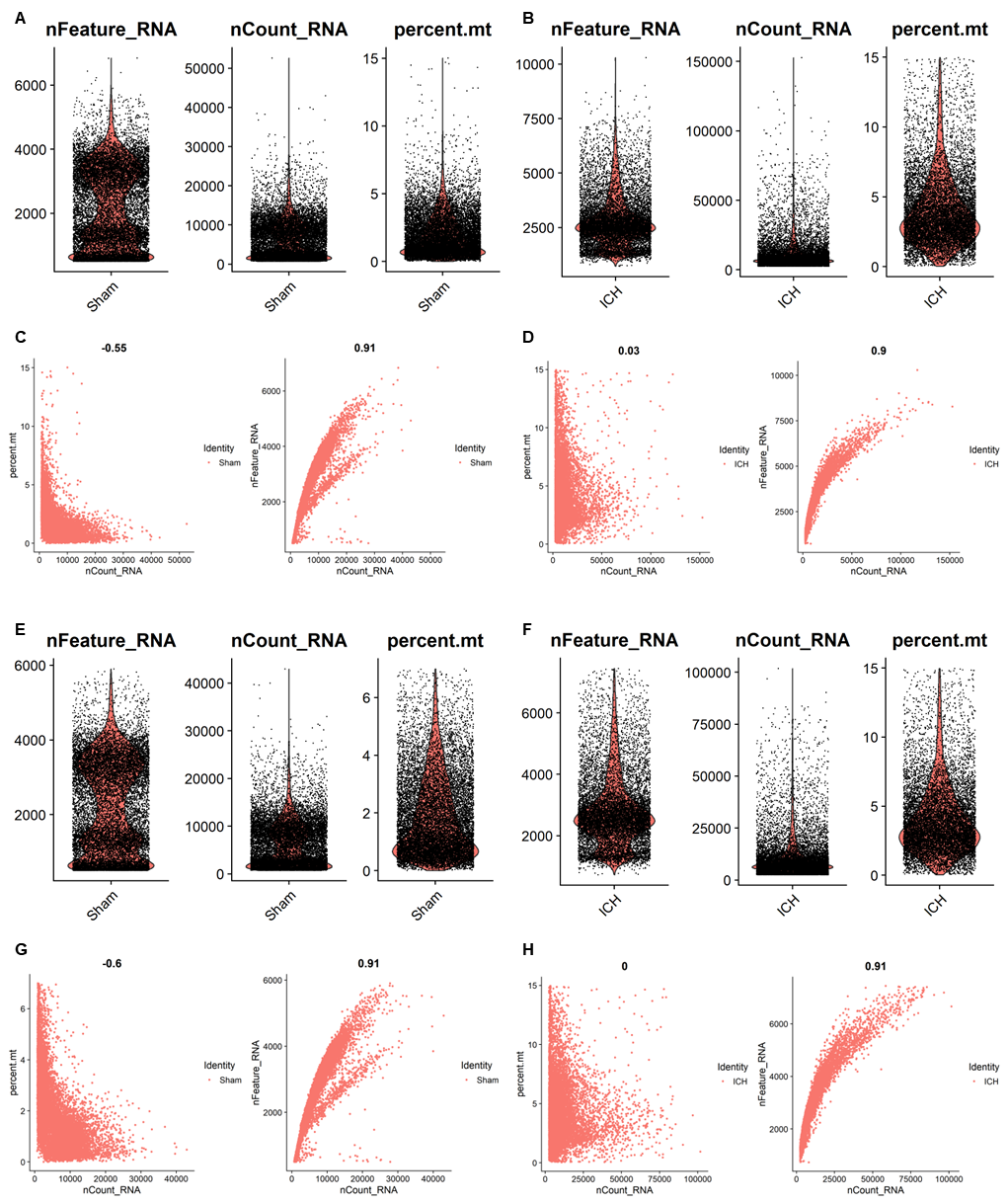

**Supplementary Figure S2.** The distributions of nFeature_RNA, nCount_RNA, and percent.mt of cells in the ICH and Sham samples before and after quality control.

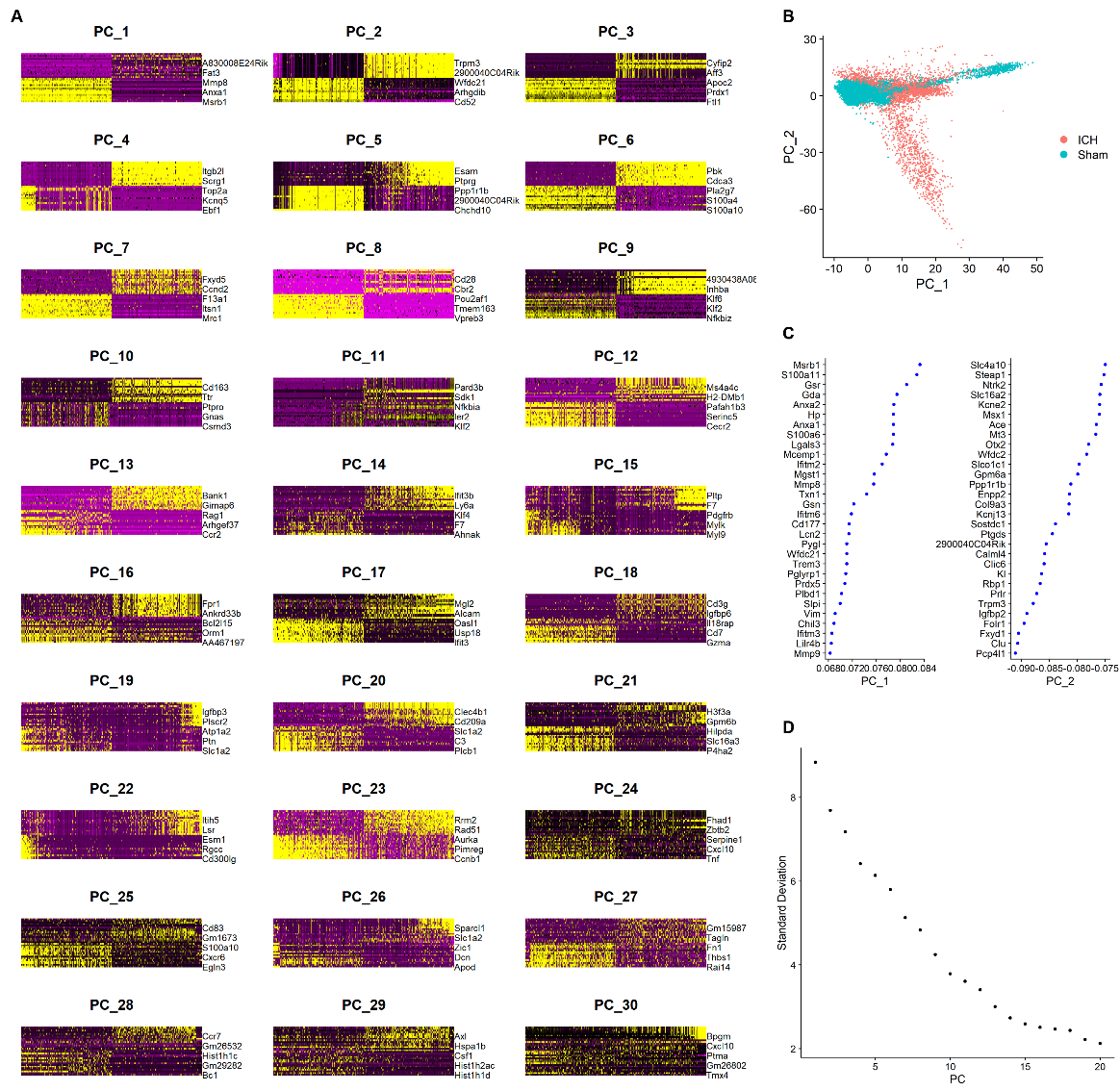
**Supplementary Figure S3. The results of principal component analysis (PCA). A.** Heatmap of the top 30 principal components. **B.** Scatter plot showing the top 2 principal components (PCs)—PC_1 and PC_2—of the ICH and Sham groups. **C.** VizDimLoadings plot showing the top contributing genes for PC_1 and PC_2. **D.** ElbowPlot displaying the standard deviation of PCs.

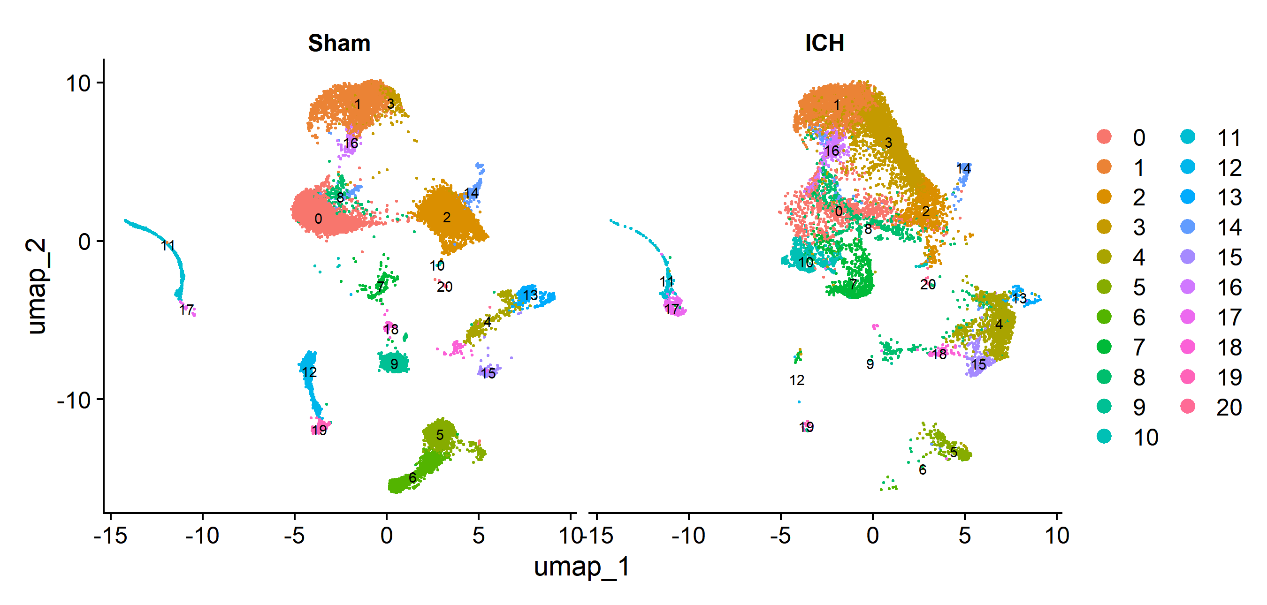
**Supplementary Figure S4. UMAP plot showing cell clustering of single-cell data in the ICH group and Sham groups.**

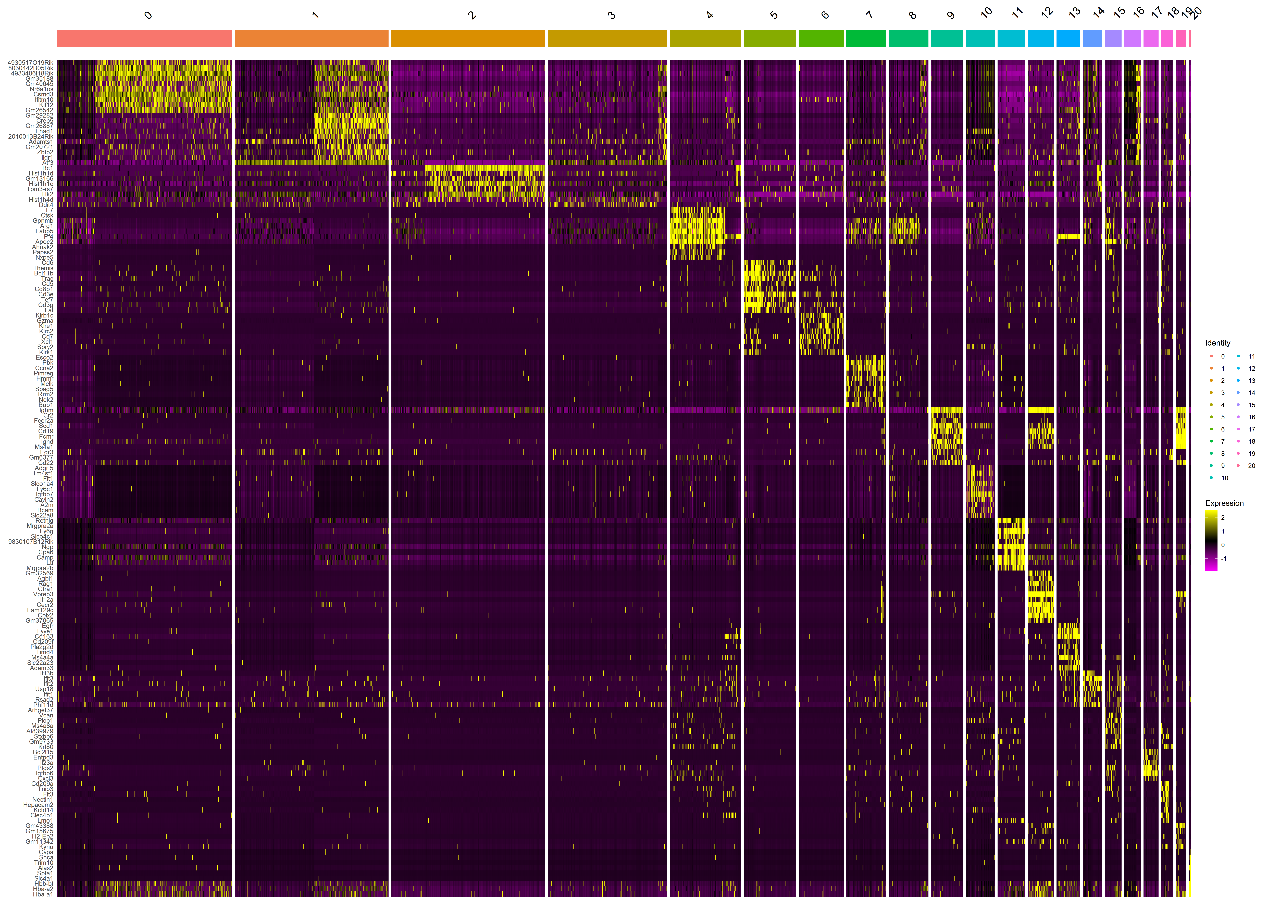

**Supplementary Figure S5. Heatmap showing the top 10 enriched marker genes of each cell cluster.**

A.

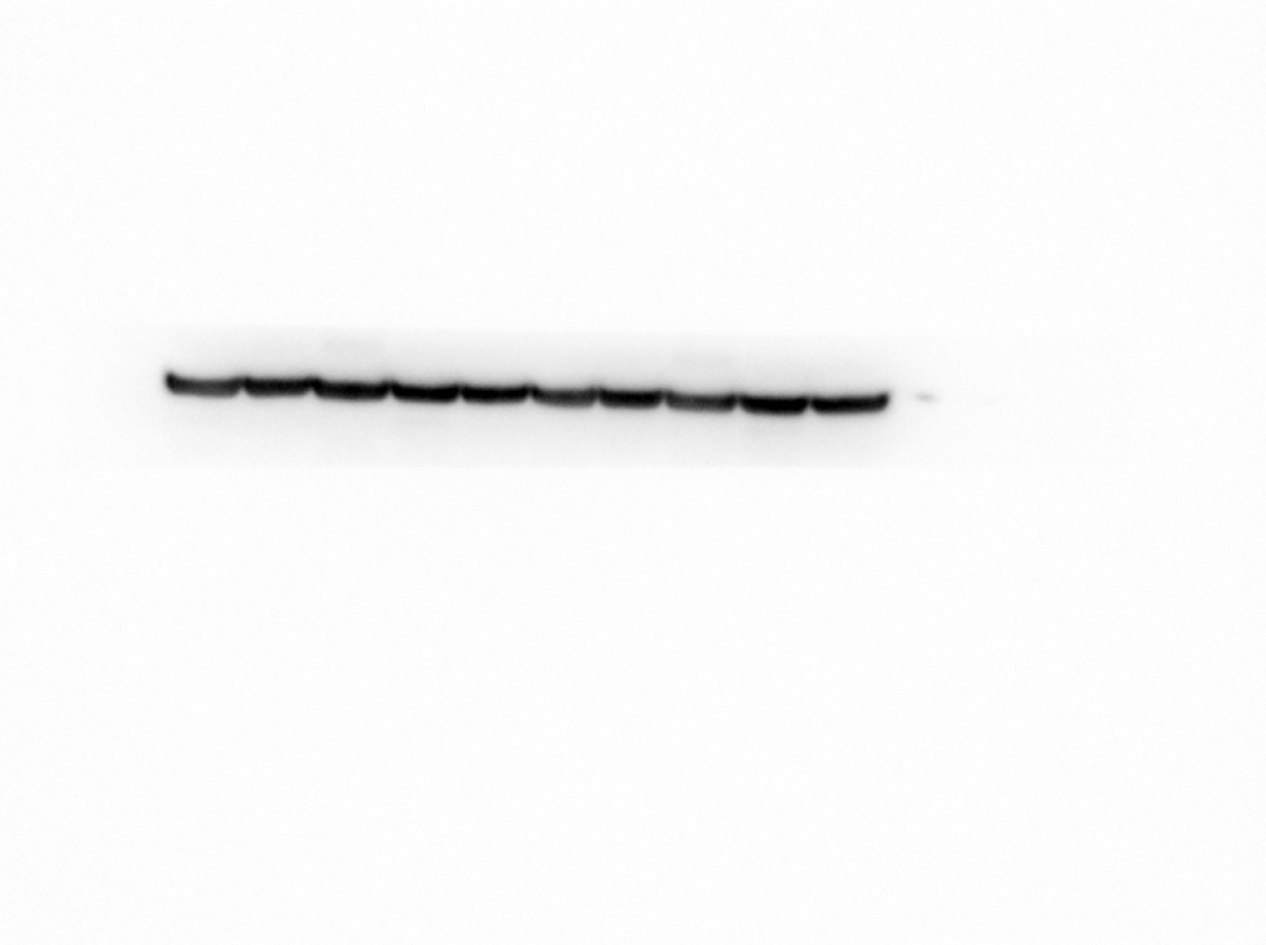

Sham 1d	3d	7d	14d

**β-actin 42 kD**

B.

**BTK 77 kD**

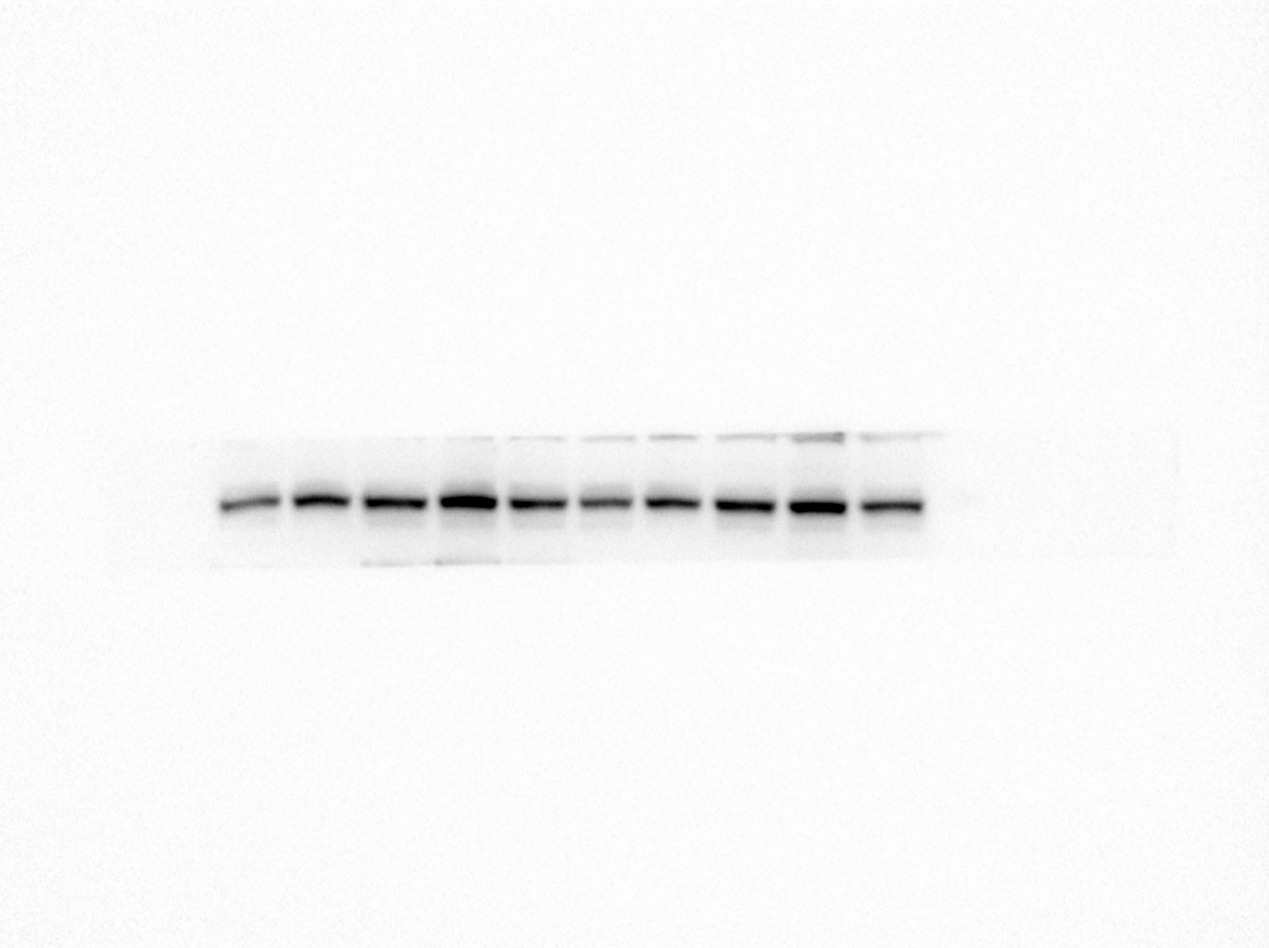

Sham 1d	3d	7d	14d

**Supplementary Figure S6. The full-length blots of Figure 2G. A.** The blot of β-actin protein. **B.** The blot of BTK protein. These two blots were from the same original large blot.

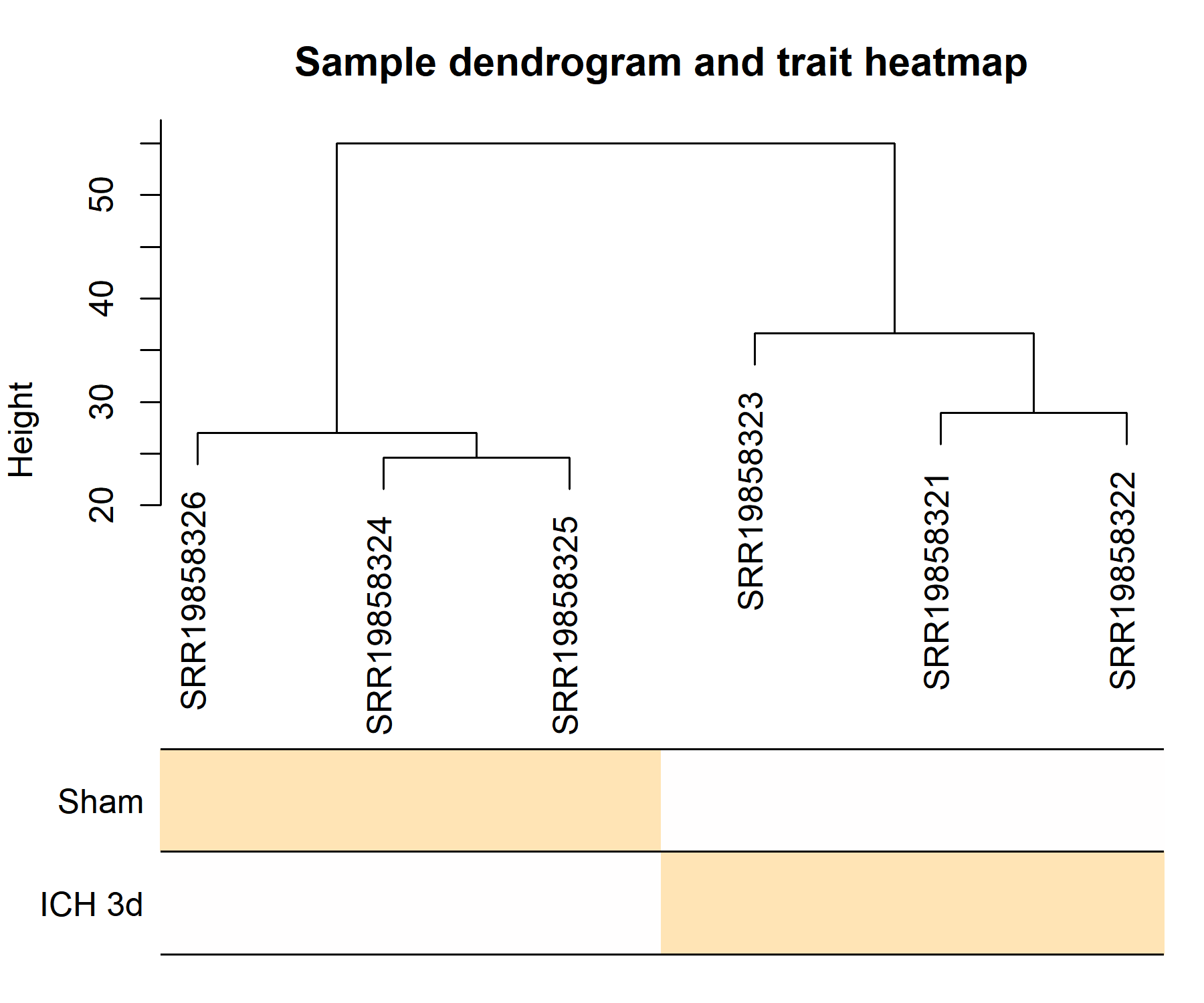
**Supplementary Figure S7.** Sample dendrogram and trait heatmap of the samples for WGCNA analysis.

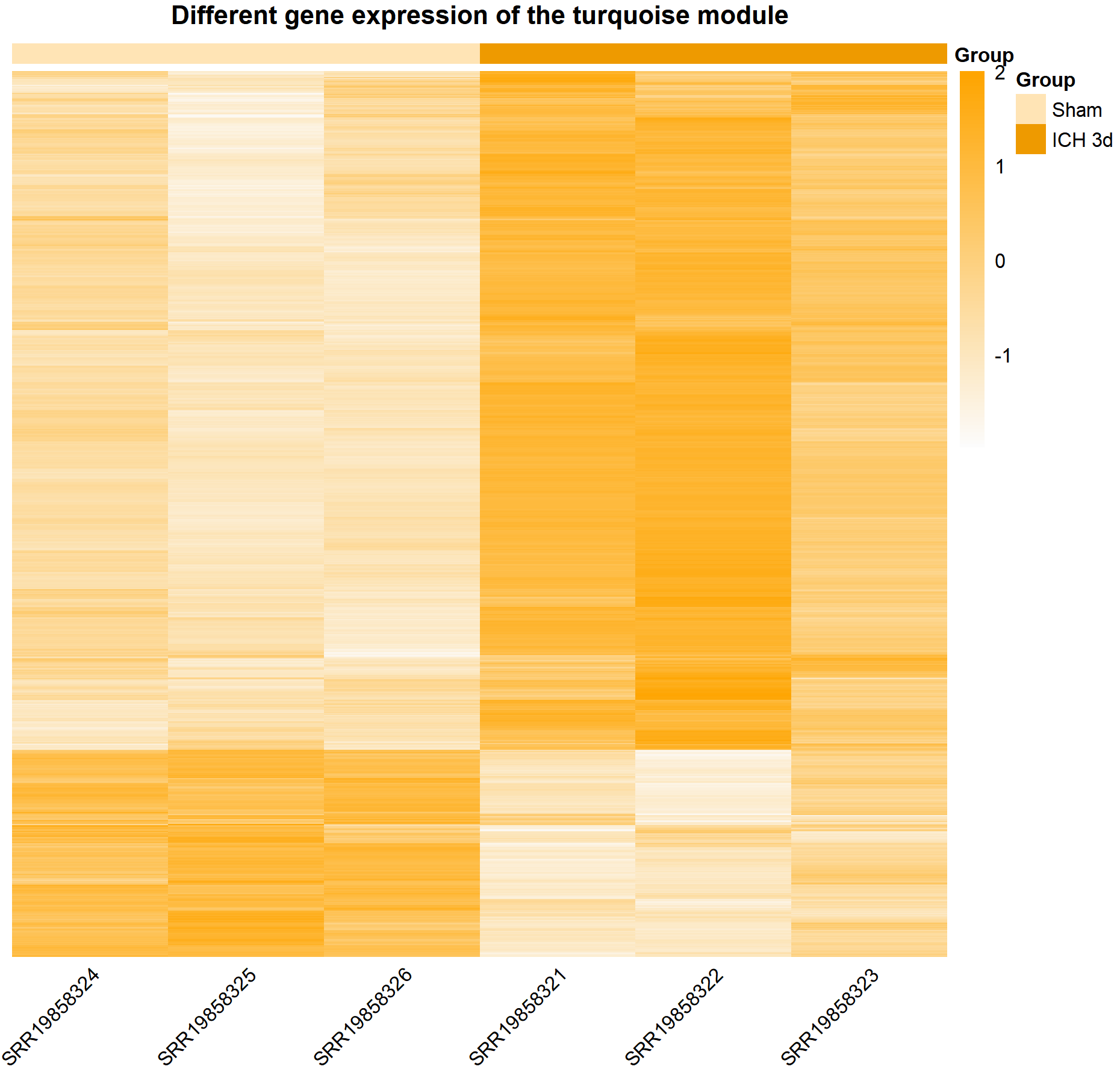

**Supplementary Figure S8. Heatmap of gene expression in the turquoise static module.**

**Supplementary Table S1. Classic cell type marker genes from the literature.**

| Ptprc | Microglia |
| --- | --- |
| Ly6g | Neutrophil |
| Gzma | NK cell |
| Ms4a1 | B cell |
| Mrc1 | M2 macrophage, M1 macrophage |
| Cx3cr1 | Microglia |
| Trbc2 | T cell |
| Cd209a | Dendritic cell |
| Il1b | Neutrophil, monocyte, M1 macrophage |

**Supplementary Table S2. The five most specifically expressed genes in each cell type.**

| Microglia | Csmd3, Nav3, Fat3, Sall3, Cacnb2 |
| --- | --- |
| Macrophage_1 | F7, Ctsk, Gpnmb, Arg1, Fabp5 |
| Macrophage_2 | Egfr, Lyve1, Cd163, Cd209f, Pla2g2d |
| Monocyte | Arhgef37, Vcan, Plcb1, Ms4a8a, AI839979 |
| Neutrophil_1 | Bcl2l15, Entpd3, Il23a, Ptgs2, Igfbp6 |
| Neutrophil_2 | Retnlg, Mrgpra2a, Ly6g, Slco4c1, Ngp |
| Dendritic cell | Cd209a, Tnip3, Tbc1d4, Flt3, Nectin1 |
| T cell | Cd6, Themis, Bcl11b, Trac, Cd5 |
| B cell | Fcmr, Ms4a1, Ly6d, Fcer2a, Cd79a |
| Innate lymphoid cell | Klrb1c, Gzma, Klre1, Klrc2, Cd7 |

**Supplementary Table S3. Marker genes used to calculate scores of microglia by AddModuleScore function.**

| **Score type** | **Marker genes** | **References** |
| --- | --- | --- |
| **Inflammatory score** | Il1a, Il18, Tnf, Tlr2, Tlr4, Nlrp3, Nfkb1, Irf5, Stat1, Socs3, Ccl2, Il6, Ccl3, Ccl4, Cxcl10 | Askenase, M. H. et al. Longitudinal transcriptomics define the stages of myeloid activation in the living human brain after intracerebral hemorrhage. Sci Immunol 6, (2021).  Langlois, J. et al. Fenebrutinib, a Bruton's tyrosine kinase inhibitor, blocks distinct human microglial signaling pathways. J Neuroinflammation 21, 276, (2024).  Zarrin, A. A., Bao, K., Lupardus, P. & Vucic, D. Kinase inhibition in autoimmunity and inflammation. Nat Rev Drug Discov 20, 39-63, (2021). |
| **M1 score** | Tnf, Il6, Il1b, Cxcl10, Nos2, Ccl4, Ccl3, Ccl12, Cd86, Fcgr2b, Fcgr1, Stat1, Nfkb1, Socs3, Fcgr3, Cd68, Il2, Il23a, Ifng, Ccl5, Cxcl1, Cxcl2, Cxcl10, Ccr1, Ccr5, Cxcr4, Stat3, Mmp9, Mmp12, Tlr2, Tlr4, Trem1, Hmgb1 | McAlpine, C. S. et al. Astrocytic interleukin-3 programs microglia and limits Alzheimer's disease. Nature 595, 701-706, (2021).  Stephens, R., Grainger, J. R., Smith, C. J. & Allan, S. M. Systemic innate myeloid responses to acute ischaemic and haemorrhagic stroke. Semin Immunopathol 45, 281-294, (2023).  Bai, Q., Xue, M. & Yong, V. W. Microglia and macrophage phenotypes in intracerebral haemorrhage injury: therapeutic opportunities. Brain13.5 143, 1297-1314, (2020).  Lan, X., Han, X., Li, Q., Yang, Q. W. & Wang, J. Modulators of microglial activation and polarization after intracerebral haemorrhage. Nat Rev Neurol 13, 420-433, (2017).  Sun, Y. et al. The dual role of microglia in intracerebral hemorrhage. Behav Brain Res 473, 115198, (2024). |
| **M2 score** | Tgfb1, Igf1, Arg1, Chil3, Cd36, Cd163, Mrc1, Msr1, Il10, Retnla |  |
| **M2a score** | Igf1, Arg1, Chil3, Mrc1, Retnla, Il10, Ccl17 |  |
| **M2b score** | Il10, Il1rn, Cd86, Il6, Fcgr1, Fcgr2b, Socs3 |  |
| **M2c score** | Il10, Cd163, Mrc1, Tgfb1, Arg1, Chil3, Retnla |  |
| **Homeostatic microglia score** | P2ry12, Tmem119, Cx3cr1, Fcrls |  |
